# Non-invasive skin topical delivery of hyaluronan

**DOI:** 10.1101/2025.04.03.646962

**Authors:** Weiwei Feng, Jiayi Ye, Shunmei Li, Shiqun Shao, Jiajia Xiang, Ying Piao, Qiuyu Wei, Jianbin Tang, Zhuxian Zhou, Youqing Shen

## Abstract

Hyaluronic acid (HA), a key glycosaminoglycan in the skin extracellular matrix, diminishes with aging, leading to dryness, wrinkles, and loss of elasticity. While subcutaneous HA injection remains the standard treatment, it is invasive and associated with complications. Here, we developed a zwitterionic polymer-based nanogel system (HA/OP) for noninvasive transdermal HA delivery. The nanogels, formed by complexing HA with poly[2-(N-oxide-N,N-diethylamino)ethyl methacrylate] (OP), exhibited pH-responsive behavior, remaining stable at weakly acidic skin surface pH (∼5) but dissociating in neutral dermal environments. HA/OP nanogels efficiently permeated the stratum corneum via sebum-assisted enrichment and para-corneocyte transport, followed by transcytosis and enhanced paracellular penetration mediated by myosin light-chain kinase (MLCK) upregulation. This enabled deep HA deposition in the dermis and subcutaneous tissue, significantly improving skin hydration and reducing wrinkles in hairless mice without irritation. Our findings present a safe, needle-free strategy for topical HA delivery, addressing a major unmet need in cosmetic and dermatological applications.

## Introduction

Hyaluronan (HA), a linear glycosaminoglycan consisting of repetitive disaccharide units of N-acetyl-D-glucosamine and D-glucuronic acid^1–3^, is a dermal filler abundant in the skin extracellular matrix^4–6^. With aging the HA content in the skin decreases gradually^7–9^, making the skin dry, rough, wrinkled, and losing plasticity^7,10,11^. Supplementing HA can smooth wrinkles simply by increasing physical volume to support the skin and provide structural support to the extracellular matrix for collagen production^12,13^. Therefore, HA as a safe and nonimmunogenic dermal filler has been used for facials, generating enormous market demand^14^.

The skin is impermeable to large macromolecules, particularly hydrophilic HA, because of its hydrophobic stratum corneum (SC)-covered multi-layered wall-like structure^15^, so subcutaneous injection of HA with needles, needleless injectors, or microneedles into the skin is still a routine procedure.^16–18^ However, the injection causes skin damage, lesions, and other severe complications.^14,19–22^ Therefore, the cosmetic industry has long dreamed of non-invasive skin topical delivery of HA by enhancing HA permeation,^23–26^ for instance, using skin-penetration peptides,^26^ but it is extremely challenging and not very successful yet^27^.

Herein, we report a polymer carrier, poly[2-(*N*-oxide-*N*,*N*-diethylamino)ethyl methacrylate] (OP), capable of complexing HA, fast permeating the skin, and depositing HA in the viable dermis and dermis, realizing non-invasive skin topical delivery of HA and cross-linked HA (cHA). OP is a pH-dependent zwitterionic polymer, neutral at pH around 7 but cationic at pH 5 or lower.^28^ So, OP and HA formed stable hybrid nanogels (named HA/OP) under weakly acidic conditions but separated quickly once in a neutral environment. On the skin surface, which is rich in fatty acids in the sebum and para-corneocyte lipids and so the pH is weakly acidic (around 5), OP bound the fatty carboxylates via electrostatic attraction, enriching HA/OP nanogels in the sebum and para-corneocyte space of the SC. Then, the nanogels overcame the SC barrier through the para-corneocyte space, and continued to penetrate through the transcytosis-mediated intracellular pathway and the upregulated MLCK-mediated paracellular permeability into the viable epidermis, dermis, and even subcutis, where the neutral microenvironment broke up the nanogels and deposited HA there, while OP further entered the circulation and was excreted by the kidneys. So, topically applying HA/OP significantly relieved the skin wrinkles and roughness of hairless SHJH hr/hr mice. Moreover, HA/OP caused no skin damage or irritation upon repeated applications. So, HA/OP nanogels provide a safe, effective, and non-invasive strategy for skin topical delivery of HA.

## Results and Discussion

### Fabrication and skin penetration of hybrid HA/OP nanogel

Zwitterionic OP was positively charged at pH 5 or lower due to the protonation of its oxygen anions (Supporting Information Fig. S1A). So, OP bound negatively charged HA with constants (K_a_) of 4.03×10^6^ mol^−1^ at pH 5, 9.43×10^6^ mol^−1^ at pH 4, and 2.28×10^7^ mol^−1^ at pH 2, as measured by isothermal titration calorimetry (ITC) (Fig. 1A). At pHs lower than 2, HA was fully protonated and could not bind OP anymore (Supporting Information Fig. S1A). Therefore, HA and OP formed complexes in a narrow acidic pH range of about 2 to 5, as further validated by aggregation-induced emission (AIE) probe (tetraphenylethylene)-modified HA (HA-TPE), whose fluorescence intensity was significantly enhanced by upon adding OP at pH 3-5 while the agglomeration and precipitation occurred due to the strong interaction force at pH 2 (Supporting Information Fig. S1B). Coarse-grained molecular dynamics (CGMD) simulations of 40 HA chains with 20 repeat units and 10 OP chains with 13 repeat units depicted the formation of the complexes (Fig. 1B). At pH 4.5 and an HA/OP mass ratio of 5, 4.8 KDa OP formed spherical NCs with a diameter of 158 nm with 5 KDa HA (Supporting Information Fig. S1C) or 207.5 nm with 200 KDa HA (Fig. 1C). The zeta-potentials of these NCs were similar, about -24 mV, and they were all stable at pH 4.5 at room temperature even for 30 days (Fig. 1C&Supporting Information Fig. S1C). Furthermore, OP could also bind crosslinked HA nanogels (cHA) to form larger hybrid NCs. For instance, 5 KDa HA was crosslinked into 322.1 nm cHA nanogels (PDI: 0.226). OP (4.8 KDa) complexed the cHA and formed cHA/OP NCs of 552.7 nm in diameter and -23.4 mV in zeta potential at pH 4.5 (Supporting Information Fig. S1D). The NCs of HA (200 KDa)/OP (4.8 KDa) were used in further research.

**Fig. 1.**
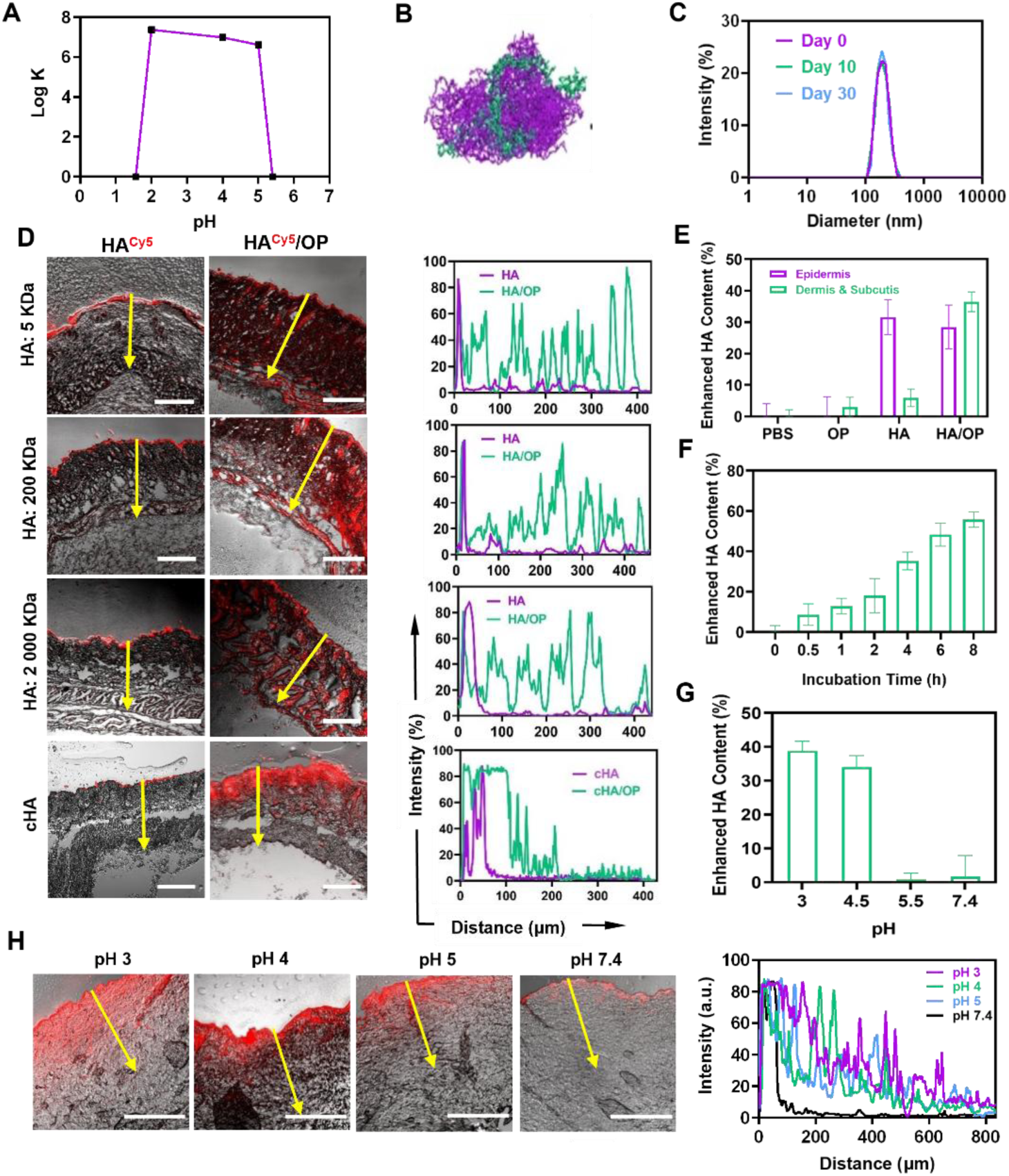
Fabrication and skin penetration of hybrid HA/OP nanocomplexes (NCs). (A) The pH-dependent binding constant of HA and OP measured by isothermal titration (HA, 1 mg/mL; HA/OP (mass) = 5). (B) A snapshot of coarse-grained molecular dynamics (CGMD) simulations at 600 ns of 40 HA (20 units, purple) and 10 OP (13 units, green). (C) The sizes of NCs of HA (200 KDa) and OP (5 KDa) at a mass ratio of 5 and pH 4.5 measured by dynamic laser light scattering at day 0, 10, and 30. (D) Representative images and Cy5-fluorescence intensity from the skin surface to deep skin layers along the randomly selected lines (yellow arrow) of the mouse skin slices after 4 h topical application with HA^Cy5^ (control) or nanogels (HA^Cy5^/OP or cHA^Cy5^/OP). (HA, 1 mg/mL; HA/OP = 5, pH 4.5). (E) The increments of the HA contents in the epidermis, and dermis and subcutis after treated with PBS, OP (0.2 mg/mL), HA (1 mg/mL), and HA/OP (HA, 1 mg/mL; HA/OP = 5) for 4 h at pH 4.5. (F, G) The incremental HA content in the dermis and subcutis of the skin treated with HA/OP (HA, 1 mg/mL; HA/OP = 5) at pH 4.5 for timed intervals (F) or 4 h at different pH values (G). In E-G, the HA contents were measured by enzyme-linked immunosorbent assay (ELISA). (H) Penetration of HA^Cy5^/OP in porcine skin at different pH: Representative CLSM images and the Cy5-fluorescence intensity from the skin surface to the deep skin layer along the randomly selected lines (yellow arrows) of the slices of the porcine skin after 4 h topical application with HA^Cy5^/OP solution at different pH (HA, 1 mg/mL; HA/OP = 5, 0.2 mL); application area, 1 cm diameter. Scale bars, 200 µm (D), and 500 µm (H).

Fluorescent cyanine5 (Cy5)-labeled HA (HA^Cy5^) and its HA^Cy5^/OP NCs at a 5/1 mass ratio were used to observe their skin permeation in Balb/C nude mice (Fig. 1D). After the topical application for 4 h, HA^Cy5^, independent of its molecular weights, all stayed on the skin surface and little fluorescence was observed beneath the SC. However, the skin topically applied with HA^Cy5^/OP NCs of 5 KDa, 200 KDa, or 2000 KDa HA had strong fluoresce in all the layers, from the SC layers to the epidermis, dermis, and subcutis, even at 400 μm away from the surface, as analyzed by linescan analysis (Fig 1D). Surprisingly, the skin permeation of the NCs was not affected by the HA molecular weights. The NCs of 2000 KDa HA permeated the skin as efficiently as those of 5 KDa HA, indicating that it was the NCs that permeated the skin rather than the HA chains. Furthermore, the 500 nm cHA/OP nanogels could also permeate the SC, mainly distributing in the dermis and some reaching subcutis. The HA/OP NCs permeation in the mouse skin including epidermis, dermis, and subcutis was further quantified by enzyme-linked immunosorbent assay (ELISA). After 4 h topical application of HA, the HA content in the epidermis increased by 31.6% compared with that of the PBS-treated skin, but the HA content in the dermis and subcutis was unchanged, indicating that HA only filled the surface wrinkles rather than permeated into the skin (Fig. 1E). In contrast, the HA content increased by 36.5% in the dermis and subcutis after the skin was topically treated with HA/OP for 4 h (HA, 1 mg/mL; HA/OP (mass) = 5, pH 4.5) while filling the skin surface wrinkles similarly. Moreover, the HA content in the dermis and subcutis increased gradually with time and reached 55.9 % increment after 8 h topical application of HA/OP (Fig. 1F). The formation and stability of HA/OP NCs were pH-dependent (Fig. 1A), as is their skin permeation. The topical application of HA/OP NCs at pH 3 or 4.5 for 4 h gave HA increment in the dermis and subcutis by 38.9%, or 34.1% relative to the original skin, while NCs at pH 5.5 or pH 7.4 could not permeate the skin (Fig. 1G), agreeable to the CLSM imaging of various pH HA^Cy5^/OP NCs treated porcine skin (Fig. 1H), in which HA^Cy5^/OP NCs also efficiently permeated the porcine skin, even reaching 800 μm away from the surface at the acidic pH3-5.

### Skin penetration mechanism of HA/OP NCs in SC

Daily application of testosterone-induced hair follicle-occluded mice were used for skin penetration assay to investigate whether the superior transdermal performance of HA/OP was attribute to the follicular pathway. The experimental results revealed that HA/OP could still penetrate the skin of hair follicle-plugged mice, suggesting that the skin permeation of HA/OP does not rely exclusively on hair follicle (Fig. 2A). Consequently, HA/OP must have broken through the SC barrier, and priority should be given to exploring the mechanism of its penetration through the SC by examining the intercorneocyte lipid matrix. At pH 4.5, HA/OP was found firmly binding on the skin sebum, and the surface immobilized with fatty acids, while HA alone or HA/OP at pH 7.4 could not (Fig 2F & Supporting Information Fig. S2C), suggesting that once applied on the skin, HA/OP bound the sebum and para-corneocyte lipids, which is composed of ceramide, cholesterol, and aliphatic acids^29,30^, via the protonated OP at the skin’s acidic pH.

**Fig. 2.**
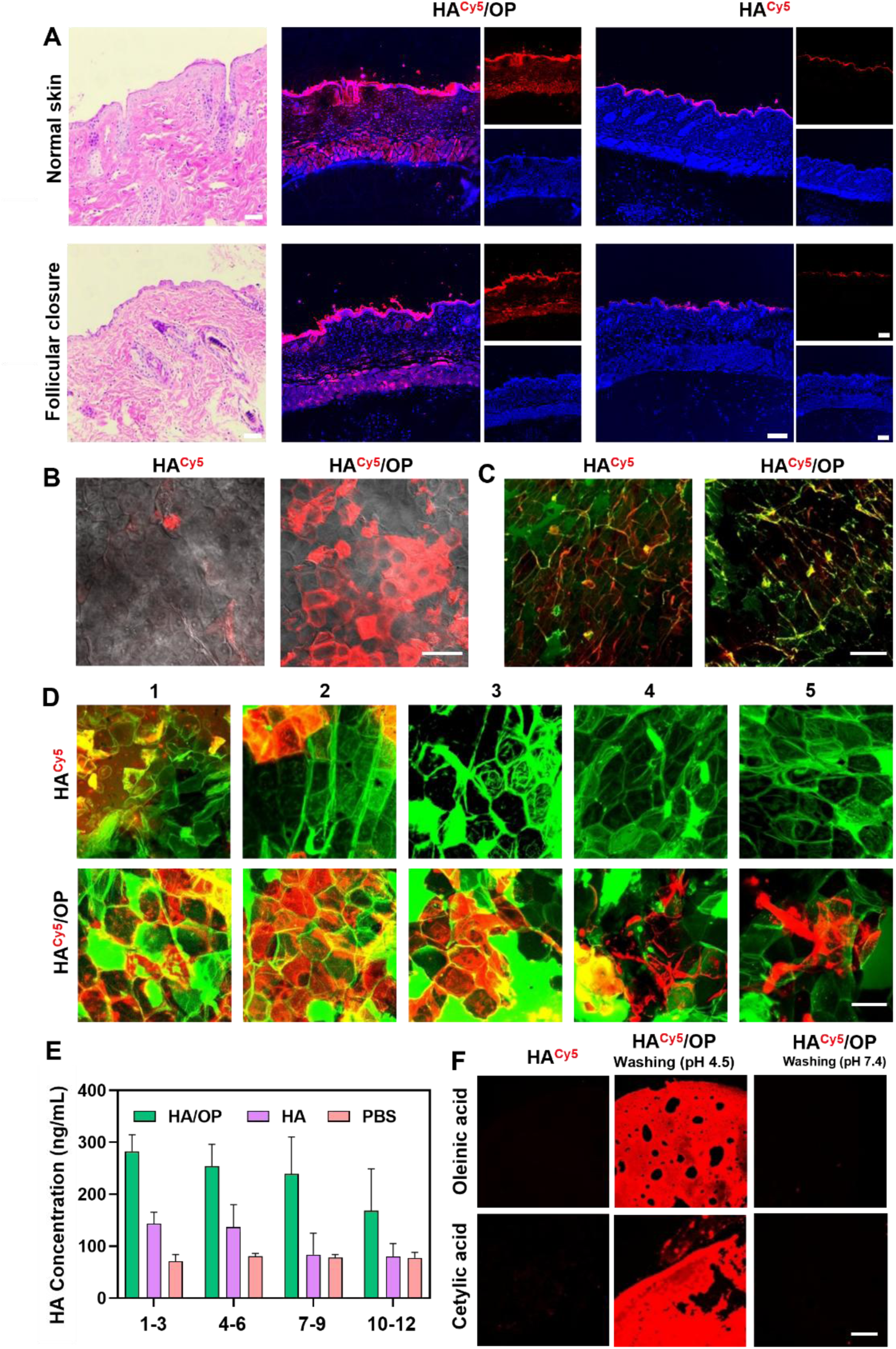
HA/OP permeation in stratum corneum. (A) Penetration of HA^Cy5^ and HA^Cy5^/OP in normal skin and hair follicle-closed skin. H&E histological analysis of normal skin and follicular-closure skin (left), the representative images of the mouse skin slices after 4 h topical application with HA^Cy5^/OP (middle) and HA^Cy5^ (right). (B) Top view of the CLSM images of the mouse ear auricles after 4 h topical application with HA^Cy5^ or HA^Cy5^/OP. (C) CLSM images of the outer SC layers on adhesive tape peeled from the rat dorsal skin after 4 h topical treatment with HA^Cy5^ or HA^Cy5^/OP. The fluorescence of Cy5 is shown in red, and the SC intercellular lipids stained with NBD-C6-HPC appear in green. (D) Fluorescence co-localization analysis of HA^Cy5^ (red) and keratinocyte space labeled with NBD-HPC-C6 (green) in the different outer SC layers on adhesive tape peeled from the mouse dorsal skin. (E) The amount of hyaluronan in the different outer SC layers on adhesive tape peeled from the mouse dorsal skin treated with HA/OP, HA, or PBS for 4h. (F) Cy5 fluorescence detection of the surface immobilized with oleinic acid and cetylic acid treated with HA^Cy5^ and HA^Cy5^/OP. All these were performed at pH 4.5 (HA, 1 mg/mL; HA/OP (mass) = 5). Scale bars, 100 µm (A), 50 µm [(B) and (C)], 20 µm (D), and 200 µm (F).

The enrichment HA/OP on the skin was further visualized by CLSM imaging of a mouse ear applied with HA^Cy5^/OP, which showed that HA^Cy5^/OP surrounded corneocytes (Fig. 2B). The images of the SC layers peeled from the treated rat dorsal skins further verified their colocalization, showing that HA^Cy5^/OP overlapped with NBD-HPC-C6-stained para-corneocyte lipids (green) and appeared yellow (Fig. 2C).

The distribution of HA in the SC layers of the mouse dorsal skin treated with HA^Cy5^ or HA^Cy5^/OP was investigated by peeling the SC layers using adhesive tape (Fig. 2D and Supporting Information Fig. S2A). Not surprisingly, HA^Cy5^-treated skin had Cy5-fluorescence limited in the two peels of the SC layers, while Cy5 fluorescence was observed in all the SC layers of the skin treated with HA^Cy5^/OP. Quantitation of the HA contents in the SC layers by ELISA confirmed the above observation, and a significant amount of HA reached the 10-12 peels of the SC layers in the HA/OP-treated skin (Fig. 2E). These results indicated that HA/OP quickly bound and were enriched on the acidic sebum and the SC outer layer, and then diffused through the para-corneocyte lipids to penetrate the SC layers.

Acidifying the skin facilitated the penetration of HA^Cy5^/OP into deeper skin. Applying HA^Cy5^/OP NCs at pH 4.5 on the skin delivered HA in the skin but did not reach the tissues under the subcutis (Supporting Information Fig. S3A). Acidifying the skin by intradermal injection of pH 3 PBS enabled the NCs to reach superficial musculo-aponeurotic system (SMAS) fascia (Supporting Information Fig. S3A & S3B). These results indicated that it was OP that carried the NCs to penetrate the skin and OP further penetrated into the tissue and entered the blood system for clearance.

### Two simultaneous pathways that HA/OP mediated in epidermis and dermis

Strong Cy5 fluorescence was also observed in the viable epidermis and dermis layers peeled from the skin treated with HA^Cy5^/OP NCs, while only sporadic weak Cy5 fluorescence could be detected in HA^Cy5^ group, which was packaged in the skin pores (Supporting Information Fig. S3D).

Therefore, it was necessary to explore the mechanism of HA/OP NCs penetration after crossing the SC to the living epidermis and dermis. The SC layers of the mouse skin were peeled off using adhesive tape (20 times), and then HA^Cy5^/OP or HA^Cy5^ at pH 4.5 or pH 7.4 was topically applied to the SC-removed area for 4 h. The skin areas were dissected and sectioned for CLSM (Fig. 3A&Supporting Information Fig. S4B), and the other parts were digested into single-cell suspensions for flow cytometry analysis (Fig. 3B&Supporting Information Fig. S4A). Without the SC, a small quantity of HA^Cy5^ could also penetrate the epidermis and dermis, giving sporadic Cy5 fluorescent spots; however, HA^Cy^^5^/OP distributed extensively in the whole skin layers even in pH 7.4 (Fig. 3A). The flow cytometry data showed that about half of the digested cells in the HA^Cy5^/OP group had HA^Cy5^, achieving the mean fluorescence three times that of the HA^Cy5^ group in pH 7.4 (Fig. 3B), which was consistent with the results under acidic conditions (Supporting Information Fig. S4A&S4B). Meanwhile, the primary keratinocytes and dermal cells isolated from whole skin treated with HA^Cy5^ or HA^Cy5^/OP was used for investigation of cellular internalization. Only with HA^Cy5^/OP, both keratinocytes and dermal cells was detected with strong Cy5 fluorescence (Supporting Information Fig. S2B). Therefore, it could be concluded that both the viable epidermis and dermis cells efficiently took up HA/OP NCs. Human keratinocytes cells (HaCaTs) were cultured into 3D spheroids to mimic epidermal tissue of the skin, and HA^Cy5^/OP was able to promote the penetration of HA^Cy5^ in HaCaT spheroids after 2 h and 4 h of incubation and bring HA^Cy5^ to the center of the spheroids at 4 h (Supporting Information Fig. S4C). The above data indicated that OP in NCs was of great significance for promoting the deep penetration of HA in epidermis and dermis.

**Fig. 3.**
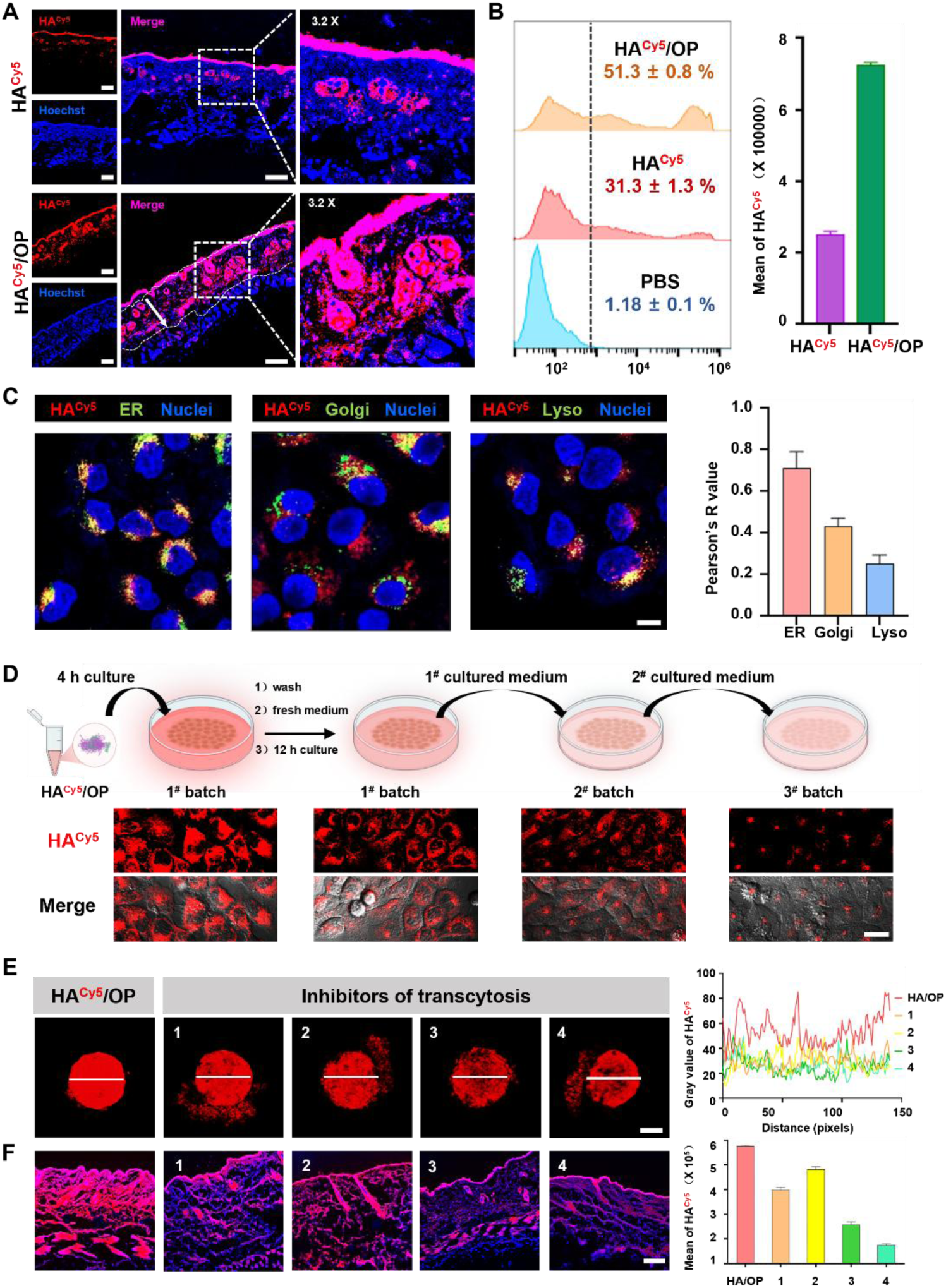
The transcytosis mechanism of HA/OP in epidermis and dermis. (**A**) CLSM images of Cy5 fluorescence in the viable epidermis and dermis of the skin, whose SC layers were removed by repeated peeling with adhesive tape 20 times and then applied with HA^Cy5^ or HA^Cy5^/OP for 4 h; HA, 1mg/mL, HA/OP (mass) = 5, pH = 7.4. (**B**) Cy5-positive cells and their main fluorescence intensity of the cells in the viable epidermis and dermis in **A**. The samples in A were digested and analyzed using flow cytometry; 10000 cells counted. (**C**) The intracellular distribution of HA^Cy5^ in organelles of HaCaT cells imaged by CLSM. The cells were cultured with HA^Cy5^/OP (HA, 0.05 mg/mL, HA/OP = 5, pH = 7.4). Cy5 fluorescence is expressed in red; ER-Tracker Green-stained ER, Golgi-Tracker Green-stained Golgi appratum, and Lyso-Tracker Green-labeled lysosomes are expressed in green, and DAPI-stained cell nuclei are in blue; Pearson number between Cy5 and each organelle’s green fluorescence was separately calculated with Fiji. (**D**) The transcytosis of HA^Cy5^/OP in HaCaT cells. The first-batch cells were cultured with HA^Cy5^/OP (HA, 0.05 mg/mL, HA/OP = 5, pH = 7.4) for 4h, and washed twice with PBS before imaging. These cells were cultured in a fresh medium for 12h, then washed and imaged; the culture medium was collected and used to culture the second batch of new cells for 12h. Furtherly, the third-batch cells were performed according to the previous treatment. (**E**) The effects of transcytosis-related inhibitors on the penetration of HA^Cy5^/OP in HaCaT spheroids and SC-removed mouse skin (**F**) imaged by CLSM (right) and analyzed by Fiji along the line (right) (1, Wortmannin; 2, Cytochalasin D; 3, Brefeldin A; 4, GW69A). The spheroids were first incubated with each inhibitor for 12 h and cultured with HA^Cy5^/OP (HA, 0.05 mg/mL, HA/OP = 5, pH = 7.4) for 4 h. The mouse dorsal skin (1 cm diameter) was peeled with adhesive tape for 20 times to remove the SC layers. Each inhibitor was subcutaneously injected into the SC-removed skin. After 1 h, HA^Cy5^ or HA^Cy5^/OP (HA, 1mg/mL, HA/OP = 5, pH = 7.4) was applied to the SC-removed area. After 4 h, the mouse was sacrificed and the skin areas were washed, dissected and sectioned into slices for imaging with CLSM (left) and (right); cell nuclei were stained with DAPI shown in blue. Scale bars, 50 μm (**A**), 10 μm (**C**), 20 μm (**D**), 100 μm [(**E**) and (**F**)].

Cell culture experiment showed that HaCaTs efficiently internalized HA^Cy5^/OP and mainly transported HA^Cy5^ into the endoplasmic reticulum (ER) and the Golgi apparatus (Fig. 3C), which are thought to be the intracellular process of transcytosis^31,32^. At the same time, experimental results of cell transmission directly proved that HA^Cy5^/OP achieved efficiently cell-to-cell delivery of HA^Cy5^ by transcytosis in HaCaTs, as evidenced by persistent Cy5 fluorescence detection up to the 3^#^ batch (Fig. 3D). Indeed, transcytosis-related inhibitors attenuated the penetration of HA^Cy5^/OP into the center of HaCaT spheroids (Fig. 3E), retained HA^Cy5^/OP on the skin surface and diminished its skin permeation abilityin vivo (Fig. 3F), further indicating that transcytosis of HA^Cy5^/OP among the viable epidermal and dermal cells played a major role of its permeation in these skin layers.

Transcytosis-related inhibitors did not completely block the penetration of HA/OP in epidermis, so it was speculated that there were other pathways, such as the intercellular pathway reported in the literature.^33,34^ To evaluate the changes in intercellular junctions, we monitored the transepithelial electrical resistance (TEER) of HaCaT monolayers cultured in Transwell inserts. The HaCaTs formed a dense monolayer after 6 days of culture, achieving an immobilly peak TEER value. Upon HA/OP treatment, a 60% reduction in TEER was detected (Fig. 4A) and a large amount of Cy5 fluorescence was detected in the outer chamber medium (Supporting Information Fig. S5A), which represented the enlarged intercellular gaps and the generation of paracellular permeability. As essential components of tight junctions (TJs) between adjacent cells in epidermis and dermis, ZO-1, occludin, and claudin-1 physically couple with the actin-based cytoskeleton through scaffold proteins, orchestrating the establishment and maintenance of cell polarity and morphology while modulating paracellular permeability properties^35^. Immunofluorescence quantification and immunoblotting confirmed the conserved total protein expression of ZO-1, occludin and claudin-1 in mouse skin after 4 h HA/OP administration (Fig. 4F), while the high-resolution imaging strikingly documented their spatial redistribution from tight junction-associated linear arrays to cytoplasmic dispersion in HaCaTs, suggesting active disassembly of junctional complexes (Fig. 4B&Supporting Information Fig. S5B). Meanwhile, actin participates in a number of critical cellular processes, including establishment and maintenance of cell junctions^36,37^. Fibros actin (F-actin) includes stress fibers, dense peripheral band (DPB), and central staple fibers, which is arranged in a filamentous and orderly manner, forming a complete and continuous actin band in normal cells as the figure shown (Fig. 4C). In the HA/OP-treated cells, the expression of F-actin increased with redistribution, which specifically referred to the rupture of the dense band of actin along the cell perimeter and the appearance of a large number of parallel bundled stress fibers near the nucleus in the center of the cell, which made the contractile force exceed the adhesion force as indicated by the arrow in the figure 4C, suggesting cell contraction and enhanced paracellular permeability (Fig. 4C). In addition, HaCaTs treated with OP alone exhibited almost unanimous changes as described above, while the fluorescence intensity and distribution of F-actin treated with HA alone were consistent with those of normal cells, indicating that the shrinking cell morphology and enhanced paracellular permeability were directly mediated by OP (Supporting Information Fig. S5C). The underlying mechanism of the alteration in TJ integrity and the formation of paracellular gap has been further explored.

**Fig. 4.**
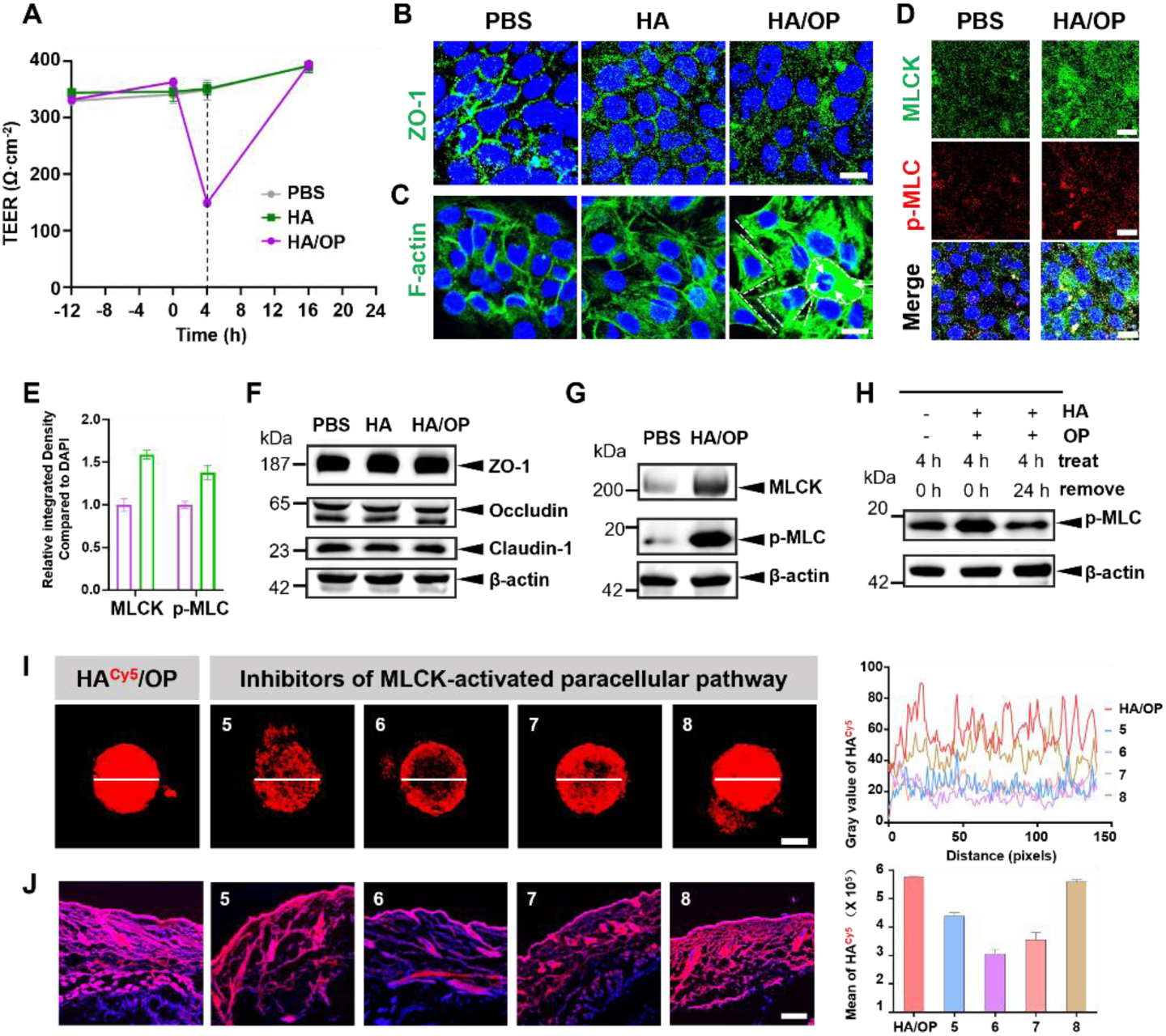
The upregulated MLCK-mediated paracellular permeability of HA/OP in epidermis and dermis. (A) Transepithelial electrical resistance (TEER) of cultured HaCaT cell monolayers treated with PBS, HA or HA/OP (HA, 0.05 mg/mL, HA/OP = 5) for 4h. TEER measurements were performed at baseline (pre-dose), and at 4 h post-dose and 12 h after the withdrawal of HA or HA/OP. (B) Expression and distribution of ZO-1 in HaCaTs pretreated with PBS, HA, or HA/OP for 4 h (HA, 0.05 mg/mL, HA/OP = 5). Antibody against ZO-1 conjugated to Alexa Fluor 488 (AF488) was used to label ZO-1 (green). (C) Expression and distribution of F-actin in HaCaTs pretreated with PBS, HA, or HA/OP for 4h (HA, 0.05 mg/mL, HA/OP = 5). Phalloidin conjugated to Alexa Fluor 555 (AF555) was used to label F-actin (green). (D) Immunofluorescence of MLCK and p-MLC in HaCaT cells pretreated with PBS or HA/OP for 4h (HA, 0.05 mg/mL, HA/OP = 5). Antibody against MLCK conjugated to AF488 and antibody against p-MLC conjugated to AF555 were used to label MLCK (green) and p-MLC (red). (E) Relative integrated density compared to DAPI in (D). (F) Western blot (WB) analysis of ZO-1, occludin, and claudin-1 in mouse skin pretreated with PBS, HA, and HA/OP (HA, 1 mg/mL, HA/OP = 5) for 4 h. In the WB analysis, ZO-1, occludin, and claudin-1 were detected using a rabbit anti-mouse primary antibody, followed by incubation with a goat anti-rabbit secondary antibody conjugated with horseradish peroxidase (HRP) for signal amplification and visualization. (G) The changes in expression of MLCK and p-MLC in mouse skin pretreated with PBS or HA/OP (HA, 1 mg/mL, HA/OP = 5) for 4h. (H) WB analysis of p-MLC in mouse skin pretreated with HA/OP (HA, 1 mg/mL, HA/OP = 5) for 4h, and p-MLC expression 24h after HA/OP removed. (I) The effects of MLCK-activated paracellular pathway on the penetration of HA^Cy5^/OP in HaCaT spheroids and SC-removed mouse skin. (J) Imaged by CLSM (left) and analyzed by Fiji along the line (right) (5, Chlorpromazine; 6, ML-7; 7, ML-9; 8, Y-27632). The spheroids were first incubated with each inhibitor for 12 h and cultured with HA^Cy5^/OP (HA, 0.05 mg/mL, HA/OP = 5, pH = 7.4) for 4 h. The mouse dorsal skin (1 cm diameter) was peeled with adhesive tape for 20 times to remove the SC layers. Each inhibitor was subcutaneously injected into the SC-removed skin. After 1 h, HA^Cy5^ or HA^Cy5^/OP (HA, 1 mg/mL, HA/OP = 5, pH = 7.4) was applied to the SC-removed area. After 4 h, the mice were sacrificed and the skin area were washed, dissected and sectioned into slices for imaging with CLSM (left) and (right); cell nuclei were stained with DAPI shown in blue. Scale bars, 20 μm [(B), (C) and (D)], 100 μm [(I) and (J)].

The phosphorylation of myosin light chain (p-MLC), which is known to mediate actin-myosin interactions, thereby regulating cell contraction and paracellular permeability ^38^. As can be seen from the confocal images, p-MLC and its one of the upstream kinase, myosin light chain kinase (MLCK) were both significantly enhanced after 4 h of HA/OP incubation in HaCaTs (Fig 4D&4E), consistent with the remarkable upregulation of p-MLC and MLCK in mouse skin showed by the western blot (WB)(Fig 4G). Inspiringly, p-MLC was recovered after 12 h of HA/OP removal in mouse skin (Fig 4H), which implied that the enhanced skin permeability resulting from the upregulated MLCK-mediated paracellular permeability mediated by HA/OP is reversible rather than permanent (Fig 4G). The amount and distribution of MLCK and ZO-1 before and after removal also verified the recovery of HA/OP-mediated paracellular permeability (Supporting Information Fig. S6A&S6B).

Consumedly attenuated fluorescence in the center of the HaCaT spheroids was the result of inhibitors associated with MLCK-p-MLC pathway (Fig 4I), while the inhibitor of Rho-associatedcoiled-coil kinase (ROCK), as another common kinase that promotes phosphorylation of myosin light chains, had no weakening effect on penetration (Supporting Information Fig. S7). Similarly, the same results were given for other endocytosis inhibitors unrelated to transcytosis, such as Filipin and Genistein (Supporting Information Fig. S7). To further validate in vivo, the TAPed-skin area topically treated by HA^Cy5^/OP was detected by FCM (Fig 4J, right) and CLSM (Fig 4J, left) after 1 h after intradermal injection of MLCK-p-MLC related inhibitors. The HA^Cy5^ fluorescence value was reduced to varying degrees, accompanied with the confined and blocked fluorescence in the outermost layer of the skin under the action of these inhibitors. However, skin treated with those inhibitors independent of two aforementioned transdermal mechanisms, such as Filipin, Genistein and Y-27632, exhibited the same superior skin penetration as the HA^Cy5^/OP positive group (Supporting Information Fig. S8).

These results indicated that the permeation effect of HA/OP in epidermis and dermis relied on two pathways that coexist, transcytosis-induced intracellular transmission and upregulated MLCK-mediated paracellular permeability in keratinocytes and epidermal cells.

### Skin-smoothing effect and biocompatibility of HA/OP

Accordingly, the wrinkle-reduction after topical skin application of HA/OP was first tested on C57BL/6 aged (18 months old) mice. Their hair was removed using Veet hair removal cream to expose the skin, which was then topically applied with HA/OP every day for 4 h. The skin became smoother and presented in a plump and satin stateon day 3 (Fig. 5A). Please note that the activation of hair follicle made skin darker gradually.

**Fig. 5.**
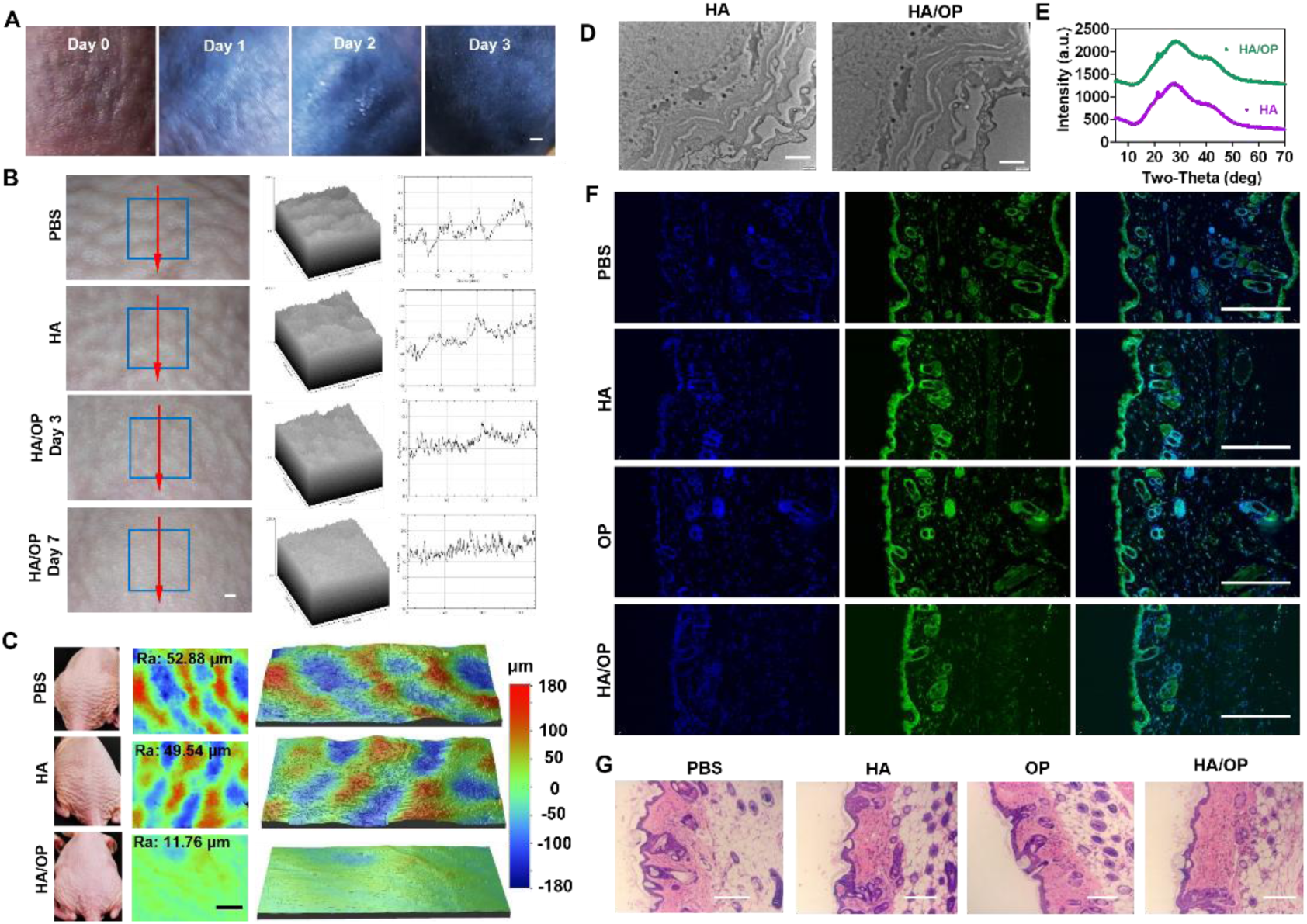
Skin-smoothing effect, and biocompatibility evaluation of HA/OP. (A) The representative images of the 18 months-aged mouse back skin treated with HA/OP once a day (HA, 1 mg/mL; HA/OP = 5; pH 4.5; 4 h); Scale bars, 1 mm. (B) The images of the SHJH hr/hr mouse skin areas treated with PBS, HA (1mg/mL), or HA/OP (HA, 1 mg/mL; HA/OP=5; pH 4.5; 4 h) once a day for 7 days (left) and SHJH hr/hr mouse skin roughness (Ra) analysis with Fiji/image J, including surface roughness by simulated images of the parts in blue frame (middle) and gray value of the position along the red lines (right). Scale bars, 1 mm. (C) The images of the SHJH hr/hr mouse skin areas treated with PBS, HA (1 mg/mL), or HA/OP (HA, 1 mg/mL; HA/OP=5; pH 4.5; 4 h) once a day for 7 days (left) and the images analyzed with a three-dimensional interference microscopy of their invert silicone models (middle and right).(D-G) Skin structural analysis: skin morphology observation by transmission electron microscopy (D), cuticular structure analysis by X-ray diffraction of the (E), immunofluorescence of nuclear factor kB (NF-kB) activation visualized by co-localized nuclear (blue) and NF-kB (green) (F), and hematoxylin and eosin (H&E) histological analysis of the treated skin areas with HA or HA/OP (G). The mouse skin was treated with PBS, HA (1 mg/mL), OP (0.2 mg/mL) or HA/OP (1 mg/mL HA, HA/OP=5) at pH 4.5 for 4 h daily for 15 days. Scale bars, 2 μm (D), 200 μm [(F) and (G)].

SHJH hr/hr mice have rhino skin-liked rough hairless skin, which is a very challenging model for reducing wrinkles^24^, which is ideal for further evidence of the superior efficacy of HA/OP. The topical applying HA every day for 7 days had no effect on the wrinkles, whose roughness was the same as that of the PBS group in the linear and planar analysis. However, the skin became exceptionally smooth after only 3 days of application of HA/OP (1 mg/ml HA, 0.5 mL for 4h per day) and even smoother after 7 days of topical application (Fig. 5B). Silicone rubber was used to invert the treated areas and the surface roughness (Ra, the calculated average between peaks and valleys) of the inverted skin was analyzed using three-dimensional interference microscopy. The 7 day-HA-treated skin had a Ra of 49.54 μm, close to that of the PBS group (52.88 μm), but HA/OP topical treatment at the same dose substantially reduced the skin RA to 11.76 μm (Fig. 5C), demonstrating an excellent wrinkle-reducing effect of HA/OP. Furthermore, as rhino skin-liked skin lack of sweat gland and hair follicle appendages, these results also suggest that the skin permeation of HA/OP was not via skin appendages.^8^

Applying HA/OP on the skin for weeks showed no skin irritation or inflammation. TEM and X-ray diffraction showed that topical application HA/OP caused no change in the SC structure (Fig. 5D&5E). The inflammation-related cytokine analysis indicated that the NF-kB pathway was not activated (Fig. 5F), and the levels of two typical inflammatory factors, TNF-α and IL-1α, were unchanged (Supporting Information Fig. S9A&S9B). The histological study confirmed that the skin tissue treated with OP or HA/OP was not different from those treated with PBS or HA (Fig. 5G); there were also no other abnormal changes in blood biochemical analysis (Supporting Information Fig. S9C&S9D). Therefore, HA/OP for wrinkle removal had good skin topical safety and systemic biosafety, which was conducive to clinical translation.

### Conclusion

Hydrophilic macromolecules such as HA are hindered by the skin barrier, especially the SC, when they enter the skin, resulting in limited application of transdermal delivery of HA in the beauty industry. Needles, needle-free injectors, and microneedles are commonly employed to surmount these limitations, which will still cause a certain degree of pain in the user, and cause locally skin damage and inflammation. There is an urgent need for a non-invasive transdermal delivery strategy. Our HA/OP hybrid nanogel system enables non-invasive transdermal delivery of hydrophilic molecules without disrupting the skin’s structure. Furthermore, it significantly reduces skin roughness, making rough skin smoother. It provides a new approach for transdermal delivery of hydrophilic, negatively charged biomacromolecules, as well as various difficult-to-transdermally-deliver small molecule drugs.

## Method

### Labelling HA with Sulfo-Cy5, and TPE

HA (400 mg) was resolved in 10 mL anhydrous formamide in 50℃, which was remixed with EDC (175 mg) and NHS (115 mg) to activate the carboxyl groups for 2h. Furtherly, ethylenediamine (60 mg) was added to the activated HA solution for introducing free amine groups onto HA for 6h to synthesize HA-NH_2_. Sulfo-Cy5-NHS ester (dissolved in DMF) was added to the HA-NH_2_ solution, stirring for 12h. Purify the conjugate via gel filtration to remove unreacted Cy5. HA-TPE was synthesized similarly except for using TPE-NHS.

### Fabrication of HA/OP NCs

HA was resolved in deionized water (1mg/mL), in which OPDEA was added dropwise to the solution until the final concentration reached 0.2 mg/mL with ultrasound. After 30min, the solution was adjusted to pH 4.5. 4.8 KDa OP formed spherical NCs with a diameter of 158 nm with 5 KDa HA or 207.5 nm with 200 KDa HA, and ζ-potential of -24mV at pH 4.5, which were measured by dynamic light scattering using Zetasizer Nano ZS (Malvern Instrument), equipped with a 4mW He-Ne laser at a wavelength of 633nm at 25 °C. At the same time, the morphology of HA/OP NCs was analyzed by TEM and Cryo-TEM.

### The interaction of HA and HA/OP with polycarboxylate surfaces

Initially, the glass slide surface was sputter-coated with gold to facilitate subsequent chemical modifications. The surface was then treated with HS-PEG8-CH2CH2COOH, which was immobilized via gold-thiol (Au-S) bonding. Following this, solutions of HA^Cy5^ (1 mg/mL, 20 μL), OP^Cy5.5^ (0.2 mg/mL, 20 μL), and a mixture of HA^Cy5^/OP (HA^Cy5^: 1 mg/mL; OP: 0.2 mg/mL, pH 4.5 or pH 7.4, 20 μL) were applied to the functionalized surface and incubated for 10 minutes. After incubation, the slides were washed three times with PBS (pH 4.5 or pH 7.4) to remove unbound materials. Finally, the fluorescence was analyzed using confocal laser scanning microscopy (CLSM).

### ITC experiments

ITC experiments were performed at 37 °C and atmospheric pressure. Twenty-three sequential injections were made for each ITC experiment. The volume of the first injection was 20 µl, and then a constant volume of OPDEA (0.2 mg/mL, 10µl per injection) was injected into a cell filled with HA solution (1 mg/mL) at various pHs. Each titration produced a reaction heat profile.

### Study on skin permeability of HA^Cy5^, and HA^Cy5^/OP

Balb/C nude mice (female, 6-weeks old) were prepared for skin permeability, ensuring no leakage on the blank skin of mice. HA^Cy5^ (1 mg/mL, 0.4 mL), HA^Cy5^/OP (HA^Cy5^, 1 mg/mL; OP, 0.2 mg/mL; pH 4.5, 0.4 mL), and cHA^Cy5^/OP (cHA^Cy5^, 1 mg/mL; OP, 0.2 mg/mL; pH 4.5, 0.4 mL) solution were injected into the glue cover that adheres to the back skin separately to ensure that the liquid completely covers the skin. After wet for 4 hours, the glue cover was removed, and the skin were washed with PBS for 3 times. Finally, the wetted skin was harvested, fixed by paraformaldehyde for 2 hours, and buried by OCT embedding agent. 10 μm thickness slices were prepared using a cryostat microtome and observed using CLSM.

Porcine skin was achieved from YSKD company in Beijing. HA^Cy5^/OP (HA^Cy5^, 1 mg/mL; OP, 0.2 mg/mL, 0.4 mL, pH 3, 4, 5, and 7.4) was prepared for further study of penetration of HA^Cy5^/OP in porcine skin at different pH. The RYJ-12B model developed by Huanghai Pharmaceutical Testing was used for detection of porcine shin permeability of HA^Cy5^/OP. PBS (pH 7.4) was added to the receptor chamber until it was full. Fix the pig skin between the donor chamber and the receptor chamber of the diffusion cell, ensuring that the skin was tightly in contact with the cell body to avoid leakage. 1 mL of HA^Cy5^/OP solution was added to the donor chamber at 37°C, while stirring device was to ensure uniform mixing of the receiving solution in the receptor chamber. The skin was harvested after 4 hours for CLSM analysis.

### ELISA qualification of HA/OP penetration in skin

The HA/OP NC permeation in the mouse skin including epidermis, dermis, and subcutis was quantified by enzyme-linked immunosorbent assay (ELISA). Balb/C nude mice (female, 6-weeks old) were prepared for qualification analysis of HA/OP skin permeability. The HA/OP NC (HA: 1 mg/mL; OP: 0.2 mg/mL, pH 4.5, 0.4 mL) was prepared according to the previous experimental protocol. A soft rubber cap was securely attached to the depilated skin of the mice to ensure no leakage. The HA solution and HA/OP solution were separately injected into the rubber cap, ensuring complete coverage of the skin. The solutions were applied for 4 h under wet conditions. After removal of the rubber cap, the skin surface was washed with PBS 3 times. Dispase II was added to the excised skin samples, followed by incubation at 37°C in a shaker for 1 h to facilitate the separation of the epidermis and dermis. Then the excised dermis was cut into small pieces. The separated tissue was minced and homogenized using RIPA strong lysis buffer. The homogenate was centrifuged, and the supernatant was collected. The HA content in the samples was quantitatively analyzed using ELISA according to the ELISA kit instruction manual.

To determine whether the penetration of HA/OP in the skin was time-dependent, we measured the HA content in the dermis and subcutaneous tissue of the wetted skin treated with HA/OP for 0.5 h, 1 h, 2 h, 4 h, 6 h, and 8 h. The acquisition of the supernatant was consistent with the method described above.

To verify whether HA/OP penetration in the skin was influenced by pH, we analyzed the HA content in the dermis and subcutaneous tissue of mouse skin treated with HA/OP at different pH values using ELISA. All the treatment was performed according to the above method except for HA/OP preparation at various pH (3, 4.5, 5.5, and 7.4).

The amount of hyaluronan in the different outer SC layers on adhesive tape peeled from the mouse dorsal skin treated with HA/OP, HA, or PBS for 4h. Balb/C nude mice (female, 6-weeks old) were prepared for qualification analysis of HA/OP skin permeability. Balb/C nude mice (female, 6-weeks old) were prepared for qualification analysis of HA/OP skin permeability in SC. The HA/OP NC (HA: 1 mg/mL; OP: 0.2 mg/mL, pH 4.5, 0.4 mL) was prepared according to the previous experimental protocol. The stratum corneum was tape-stripped 12 times, and each tape was soaked in 200 μL of deionized water for 1 hour. The collected solutions were then pooled every three tapes, and the hyaluronic acid (HA) content was quantified by ELISA.

### Skin-smoothing effect of HA/OP

C57BL/6 aged (18 months old) mice were used for skin-smoothing study. Their hair was removed using Veet hair removal cream to expose the skin, which was then topically applied with HA/OP (HA, 1 mg/mL; OP, 0.2 mg/mL, pH 4.5, 0.4 mL) every day for 4 h. Images were captured with iPhone 14 Pro rear camera every day.

SHJH hr/hr mice (16 weeks old) with rhino skin-liked rough hairless skin were used for reducing wrinkles, which was topically applied with PBS, HA (1 mg/mL), and HA/OP (HA, 1 mg/mL; OP, 0.2 mg/mL, pH 4.5, 0.4 mL) every day for 4 h. Surface morphological features of the skin were captured using a stereo microscope (Chengrixinguang, SZM-45B1), which was further analyzed with ImageJ. Meanwhile, silicone rubber was prepared for skin surface roughness analysis using 3D interference microscopy (Veeco, 9100).

### Evaluation of Biocompatibility of HA/OP

The biocompatibility of HA/OP was investigated through the analysis of inflammatory factor expression and histological examination in the skin. Balb/C nude mice (female, 6-weeks old) were subjected to daily topical application of PBS, HA, OP, and HA/OP for 15 consecutive days. After the treatment period, the treated skin was excised, embedded in OCT compound, and sectioned into 10 µm slices using a cryostat microtome. Sections were processed according to the instructions of the NF-κB Activation-Nuclear Translocation Assay Kit. Briefly, sections were fixed with fixative for 10 minutes, followed by three washes with washing buffer (5 minutes each). Immunostaining blocking solution was applied and incubated at room temperature for 1 hour. After removing the blocking solution, sections were incubated with NF-κB p65 mouse monoclonal antibody at room temperature for 1 hour, washed three times, and then incubated with anti-mouse Cy3 for 1 hour. After washing twice, nuclei were stained with DAPI for 5 minutes. Finally, sections were washed three times, mounted with anti-fade mounting medium, and observed under a confocal laser scanning microscope.

Balb/C mice (female, 6-weeks old) were subjected to daily topical application of PBS, HA, OP, and HA/OP for 15 consecutive days. After the treatment period, the treated skin was harvest. The expression of TNF-α in the treated skin tissues was quantified using a mouse TNF-α ELISA kit. Skin tissues treated with PBS, HA, OP, and HA/OP (daily for 15 days) were homogenized in RIPA lysis buffer, and the supernatant was collected after centrifugation. The ELISA procedure was performed according to the ELISA kit instruction manual.

The expression of IL-1α in the treated skin tissues was quantified using a mouse IL-1α ELISA kit. Skin tissues treated with PBS, HA, OP, and HA/OP (daily for 15 days) were homogenized in RIPA lysis buffer, and the supernatant was collected after centrifugation. The ELISA procedure was performed according to the ELISA kit instruction manual.

Hematoxylin and eosin (H&E) staining of Treated skin tissues were fixed overnight in Bouin’s fixative, embedded in paraffin, and sectioned into 5 µm slices. Sections were deparaffinized with xylene, dehydrated with 100%, 95%, and 70% ethanol, and rinsed with water. Hematoxylin and eosin (H&E) staining was performed, and images were acquired using an optical microscope.

Applying HA/OP on the skin for weeks did not show any showed no skin irritation or inflammation. TEM and X-ray diffraction showed that topical applying application HA/OP caused no change in the SC structure (Fig. 2D&2E).

Serum biochemical analysis was performed for evaluation of biocompatibility of HA/OP. Balb/C mice (female, 6-weeks old) were subjected to daily topical application of PBS, HA, OP, and HA/OP for 30 consecutive days. Blood samples were collected for serum separation by centrifugation at 3,000 × g for 15 min at 4°C. Roche Cobas 8000 automated analyzer (Roche Diagnostics, Germany) was used following the manufacturer’s protocols.

### HA/OP permeation in stratum corneum (SC)

Balb/C nude mice (female, 6-weeks old) were prepared for mice ear auricles penetration of HA^Cy^5/OP. HA^Cy5^ (1mg/mL, 0.4 mL), and HA^Cy5^/OP (HA^Cy5^: 1 mg/mL; OP: 0.2 mg/mL, pH 4.5, 0.4 mL) were injected into the glue cover that adheres to the ear auricles separately to ensure that the liquid completely covers the ear. After wet for 4 hours, the glue cover was removed, and were washed with PBS for 3 times. Finally, the wetted ear auricles were harvested for CLSM detection. Rats (female, 12-weeks old) were prepared exposing the skin via their hair was removed using Veet hair removal cream. HA^Cy5^ (1mg/mL, 0.4 mL), and HA^Cy5^/OP (HA^Cy5^: 1 mg/mL; OP: 0.2 mg/mL, pH 4.5, 0.4 mL) were injected into the glue cover that adheres to the skin separately to ensure that the liquid completely covers it. After wet for 4 hours, the glue cover was removed, and were washed with PBS for 3 times. The SC was stripped with D-square stripping disks for five times. CLSM images of the outer SC layers on adhesive tape peeled from the rat dorsal skin after 4 h topical treatment with HA^Cy5^ or HA^Cy5^/OP.

Balb/C nude mice (female, 6-weeks old) were prepared for HA/OP permeation in SC. HA^Cy5^ (1mg/mL, 0.4 mL), and HA^Cy5^/OP (HA^Cy5^: 1 mg/mL; OP: 0.2 mg/mL, pH 4.5, 0.4 mL) were injected into the glue cover that adheres to the skin separately to ensure that the liquid completely covers it. After wet for 4 hours, the glue cover was removed, and were washed with PBS for 3 times. The SC was stripped with D-square stripping disks for five times. CLSM images of the outer SC layers on adhesive tape peeled from the rat dorsal skin after 4 h topical treatment with HA^Cy5^ or HA^Cy5^/OP. Balb/C nude mice (female, 6-weeks old) were prepared for HA/OP permeation in SC. HA (1mg/mL, 0.4 mL), and HA/OP (HA: 1 mg/mL; OP: 0.2 mg/mL, pH 4.5, 0.4 mL) were injected into the glue cover that adheres to the skin separately to ensure that the liquid completely covers it. After wet for 4 hours, the glue cover was removed, and were washed with PBS for 3 times. The SC was stripped with D-square stripping disks for 12 times. Each disk was immersed in 0.2 mL PBS for 10 min, in which PBS was collected for HA ELISA analysis. The HA content in the samples was quantitatively analyzed using ELISA according to the ELISA kit instruction manual.

### Interaction of HA^Cy5^/OP with interkeratin lipids

The interaction between HA^Cy5^/OP and lipids mimicking the stratum corneum intercellular lipids, including artificial sebum, oleic acid, and cetylic acid, was investigated. Initially, the surface of a glass slide was coated with gold, and 16-mercaptohexadecanoic acid was immobilized onto the gold surface via a thiol-gold bond. Subsequently, artificial sebum, oleic acid, and cetylic acid (1 mg/mL, 100 µL) were applied onto the surface for 10 minutes, followed by washing with ethanol three times to remove unbound lipids. Then, HA^Cy5^ (1 mg/mL, 20 µL) and HA^Cy5^/OP (HA^Cy5^: 1 mg/mL; OP: 0.2 mg/mL, pH 4.5, 20 µL) were added to the treated surface for 10 minutes. Finally, the surface was rinsed three times with PBS (pH 4.5 or pH 7.4), and fluorescence was analyzed using confocal laser scanning microscopy (CLSM).

### Primary cell isolation

Balb/C nude mice (female, 6-weeks old) were prepared for skin permeability, ensuring no leakage on the blank skin of mice. HA^Cy5^ (1mg/mL, 0.4 mL), and HA^Cy5^/OP (HA^Cy5^: 1 mg/mL; OP: 0.2 mg/mL, pH 4.5, 0.4 mL) were injected into the glue cover that adheres to the back skin separately to ensure that the liquid completely covers the skin. After wet for 4 hours, the glue cover was removed, and the skin were washed with PBS for 3 times. Mice were subjected to disinfection via immersion in 75% ethanol for 5 minutes. Then, the wetted skin was harvested without excess subcutaneous tissue, and dispase II was added to the excised skin samples, followed by incubation at 37°C in a shaker for 1 h to facilitate the separation of the epidermis and dermis. The minced epidermal tissue was digested with 0.25% trypsin at 37°C for 15-20 minutes. The digested tissue was pipetted several times to dissociate the cells, and the floating epidermis was aspirated. The sample was then centrifuged at 1000 rpm for 5 minutes. After removing the supernatant, DMEM medium with 10% fetal bovine serum was added.

Then the excised dermis was cut into small pieces. The minced dermal tissue was digested with 0.2% collagenase at 37°C for 30 minutes. The digested tissue was repeatedly pipetted to dissociate the cells and then filtered through a 200-mesh sieve to remove excess tissue. The dermal cells were collected by centrifugation at 1000 rpm for 5 minutes. After removing the supernatant, DMEM medium supplemented with 20% fetal bovine serum was added. The cells were thoroughly mixed by pipetting, counted, and seeded after adjusting the cell concentration.

The separated tissue was minced and homogenized using RIPA strong lysis buffer. The homogenate was centrifuged, and the supernatant was collected. The HA content in the samples was quantitatively analyzed using ELISA according to the ELISA kit instruction manual.

### Hair follicle closure experiment

Prepared testosterone at a concentration of 5 mg/mL with a 1:1 volume ratio of a mixture of absolute ethanol and PBS. After 3 weeks of continuous application on the back of the dehaired mice, the skin tissue was stained for HE staining to determine that the hair follicle has been completely occluded. HA^Cy5^/OP or HA^Cy5^ was topically treated on the skin with enclosed hair follicles for 4 h, and then the treated skin was collected with removed subcutaneous fat and placed in 4% paraformaldehyde for fixation. Skin tissue sections with a thickness of 10 μM were obtained by embedding and ice cutting for CLSM detection.

### HA/OP permeation in SC-removed skin by CLSM and flow cytometry

Balb/C mice (female, 6-weeks old) were continuously torn off the tape until their skin was reddened and the tissue fluid was oozing, which was considered to be SC-removed. Those SC-removed mice were prepared for exploring the penetration mechanism of HA^Cy5^/OP in epidermis and dermis. HA^Cy5^ (1 mg/mL, 0.4 mL), and HA^Cy5^/OP (HA^Cy5^: 1 mg/mL; OP: 0.2 mg/mL, pH 4.5, 0.4 mL) were injected into the glue cover that adheres to the SC-removed back skin separately to ensure that the liquid completely covers the skin. After 4 h incubation, the glue cover was removed. The treated skin were washed with PBS for 3 times and cut along the inner diameter of the glue cover. Half of the skin was fixed with 4% paraformaldehyde and prepared into 10 μm thick frozen sections for Cy5 fluorescence detection, and the other half was digested into a single-cell suspension for flow cytometry. The specific digestion steps are as follows, the skin was chopped and placed into 3 mL of RPMI containing 0.25 mg/mL of Liberace and 1 μg/mL of DNase on a 37 °C shaker for 2 h. After the final 10 min incubation with the addition of 600 μL of 0.25% trypsin-1 mM EDTA, an equal volume of PBS is added to stop the enzymatic reaction. Filtered with the piston core rod of a 5 mL syringe on a sterile 100 nm membrane while grinding, collected it into a 50 mL centrifuge tube, and centrifuged at 2,000 rpm for 10 min. After resuspension with PBS, the skin single-cell suspension was obtained for flow cytometry (FCM)

For the effect of inhibitors on epidermal penetration, we injected a specific amount of various inhibitors subcutaneously before normal dosing (Wortmannin, 1.5 mg/kg; Cytochalasin D, 1.5 mg/kg; Filipin, 1.5 mg/kg; Genistein, 5 mg/kg; Chlorpromazine, 2.5 mg/kg; Brefeldin A, 5 mg/kg; GW69A, 2.5 mg/kg; ML-7, 2.5 mg/kg; ML-9, 2.5 mg/kg; Y-27632, 5 mg/kg). After an hour, the stratum corneum was removed and was topically treated at the site of the subcutaneous injection with HA^Cy5^/OP for 4 h. The rest of the procedure was the same as described above.

### Characterization of proteins involved in intercellular pathways by WB and CLSM

Balb/C mice (female, 6-weeks old) were prepared for the characterization of proteins involved in intercellular pathways by WB. ZO-1, Occludin, Claudin, MLCK and p-MLC were detected using a rabbit anti-mouse primary antibody, followed by incubation with a goat anti-rabbit secondary antibody conjugated with horseradish peroxidase (HRP) for signal amplification and visualization.

### Cultivation and penetration experiments of HaCaT 3D spheroids

3D-cultured multilayer HaCaT spheroids were prepared using the hanging-drop method. First, counted HaCaTs were resuspended with DMEM medium and mixed with sodium methylcellulose (1.2% w/v in PBS) at a volume ratio of 4:1 to a final density of 5.0 x 10^5^ cells/mL. Dropped 25 μL of the above mixture onto the inverted lid at spatial intervals and quickly covered the lid back to the cell culture plate containing PBS. After 72 h of incubation, the multilayer HaCaT spheroids were initially formed. At this point, 80 μL of agarose (1.0% w/v in PBS) was added to the 96-well plate, and the HaCaT spheroids were transferred to it with one spheroid per well for another 72 h of incubation.

The matured HaCaT spheroids were then incubated with HA^Cy5^ and HA^Cy5^/OP at a HA-equivalent dose of 25 μg/mL for 2 h or 4 h. After washing with PBS, the Cy5 fluorescence of HaCaT spheroids were detected with CLSM layer by layer from the top to the maximum diameter at 20 μm intervals. In order to explore the effects of various inhibitors on the penetration of HA/OP into the skin of the living epidermal layer, the matured HaCaT spheroids were separately pretreated with PBS, wortmannin (5 μM), cytochalasin D (5 μM), brefeldin A (10 μM), GW69A (20 μM), Chlorpromazine (50 μM), ML-7 (20 μM), ML-9 (20 μM), or Y-27632 (100 μM) for 12 h, and then replaced inhibitors with the medium containing HA^Cy5^/OP and continued to incubate for 4 h. Finally, CLSM was used to detect the Cy5 fluorescence of HaCaT spheroids treated with different inhibitors to explore the transdermal mechanism in epidermis and dermis.

### Transcytosis of HA^Cy5^/OP in HaCaT cells

HaCaTs were seeded in a confocal dish (1^#^ batch) at a density of 1.0 x 10^5^ cells/mL and incubated overnight. After 4 h of HA^Cy5^/OP treatment (at HA^Cy5^ dose of 25 μg/mL), the cells were washed with PBS and added in fresh medium for 12 h. The 12 h-cultured medium from 1^#^ batch was collected and used to treat the 2^#^ batch for 4 h, which could be understood as transcytosis-mediated HA^Cy5^ delivery by HaCaTs in the first generation. Similarly, added the medium after 12 h of exocytosis in the 2^#^ batch to the 3^#^ batch and incubated for 4 h.

### Organelle colocalization of HA^Cy5^/OP in HaCaT cells

HaCaTs were first plated at a density of 1.0 x 10^5^ cells/mL in three confocal dishes, incubated for 24 h and treated with HA^Cy5^/OP for 4 h at HA^Cy5^ dose of 25 μg/mL. Two of the dishes were added with 1 μM of Endoplasmic Reticulum-Tracker-Green dye and 0.5 μM of Lysotracker-Green dye, respectively, and incubated at 37 °C for 0.5 h. The remaining dish was washed with PBS and then added 200 μl of 5 μM Golgi-Tracker-Green dye prepared with HBSS containing 0.34 mg/mL BSA for 0.5 h incubation at 4 °C. After the organelle staining was completed, 2 μM of Hoechst nuclear dye was added and incubated for 0.5 h in a 37 °C incubator. The colocalization of HA^Cy5^ with each organelle was detected by CLSM and quantified with Pearson’s R value by Image J.

### Western Blot of TJ proteins, MLCK and p-MLC

The dehaired-skin was topically treated by HA/OP, HA, OP, or PBS for 4 h. The skin from each group was cut off and added to 500 μL of RIPA lysate buffer containing 1% protease inhibitor (PMSF) and phosphatase inhibitor. Repeatedly cut the skin with scissors for 0.5 h and then placed it on ice for 1 h. After centrifugation at 12,000 rpm for 20 min at 4 °C, the supernatant was mixed with 5X loading buffer and boiled at 100 °C for 10 min in a metal bath to complete the preparation of protein samples for subsequent use. The protein sample was loaded and the targeted proteins (ZO-1, Occludin, Claudin-1, MLCK, p-MLC and β-actin) were transferred from the gel to a membrane using a transfer buffer after electrophoresis. The expression of the target proteins was semi-quantitatively analyzed by Image J software.

### Distribution of F-actin in HaCaT cells

HaCaTs were densely plated in confocal dishes at a density of 1.0 x 10^5^ cells/mL and incubated for 24 h. After4 h of incubation with HA/OP, HA, OP, or PBS, the cells were washed with PBS, fixed with 4% paraformaldehyde for 10 min and then permeabilized with 0.5% Triton-X for 5 min at room temperature. Finally, the cells were incubated with 200 μL of PBS containing 100 nM of TRITC-labeled phalloidin for 0.5 h and stained with 200 μL of DAPI before fluorescence observation with CLSM. Selected TRITC excitation/emission filters (Ex/Em=545/570 nm) and DAPI excitation/emission filters (Ex/Em=364/454 nm).

---

**Fig. S1.**
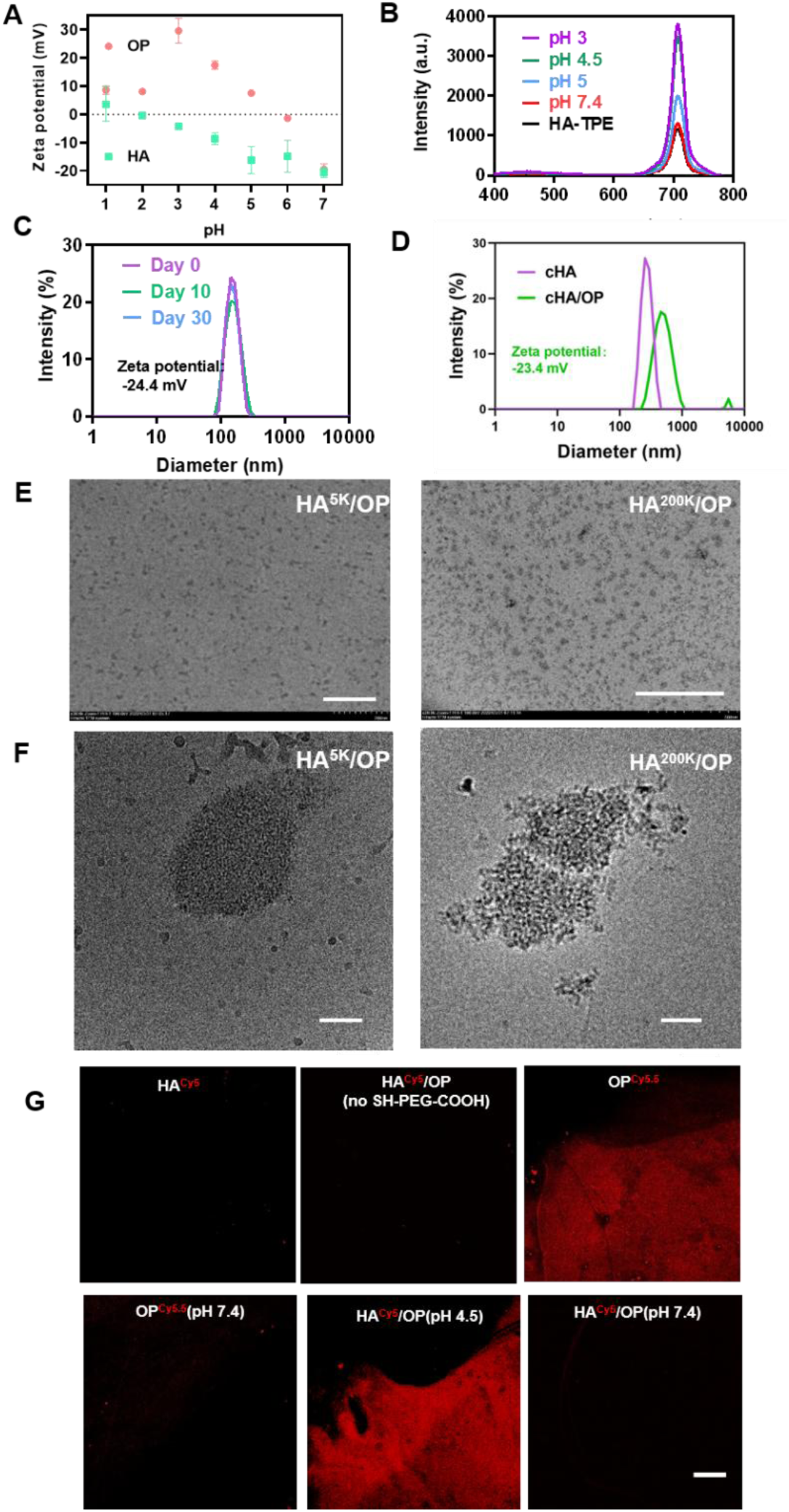
Fabrication of HA/OP. (A) Zeta potentials of HA (1 mg/mL) and OP (0.2 mg/mL) at various pHs. (B) Fluorescence intensity of HA-TPE and HA-TPE/OP in various pHs. 1mg/mL HA-TPE, HA-TPE/OP (mass) = 5. (C) The sizes of NCs of HA (5 KDa) and OP (5 KDa) at a mass ratio of 5 and pH 4.5 measured by dynamic laser light scattering at day 0, 10, and 30. (D) The sizes of NCs of cHA and cHA/OP at a mass ratio of 5 and pH 4.5 measured by dynamic laser light scattering. (E)Transmission electron microscope (TEM) graph of HA^5K^/OP (left) and HA^200K^/OP. (F) Cyro-EM of HA^5K^/OP (left) and HA^200K^/OP (right) nanogel. (G) Cy5 and Cy5.5 fluorescence detection of the surface immobilized with SH-PEG-COOH treated with HA^Cy5^, OP^Cy5.5^, and HA^Cy5^/OP. Scale bars, 200 nm (E, HA^5K^/OP), 500 nm (E, HA^200K^/OP), 100 nm (F), and 200 µm (G).

**Fig. S2.**
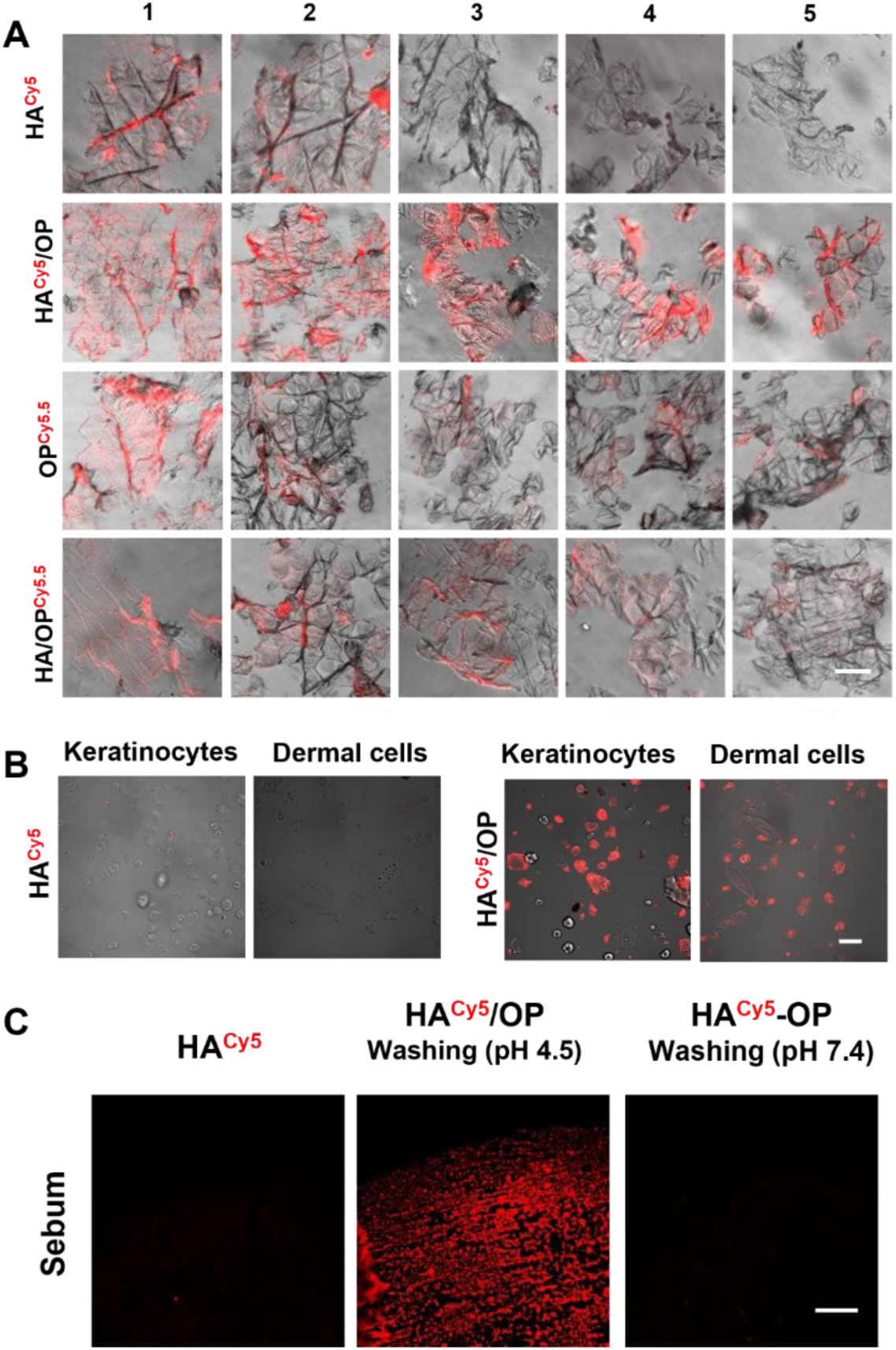
HA/OP permeation in stratum corneum. (A) Fluorescence of HACy5 (red) in the different outer SC layers on adhesive tape peeled from the mouse dorsal skin pretreated with HA^Cy5^, HA^Cy5^/OP, OP^Cy5.5^, and HA/OP^Cy5.5^. (B) Primary keratinocytes and dermal cells isolated from mouse skin predated with HA^Cy5^ and HA^Cy5^/OP for 4 h. (C) Cy5 fluorescence detection of the surface immobilized with sebum treated with HA^Cy5^ and HA^Cy5^/OP. Scale bars, 50 μm [(A) and (B)], 200 μm (C).

**Fig. S3.**
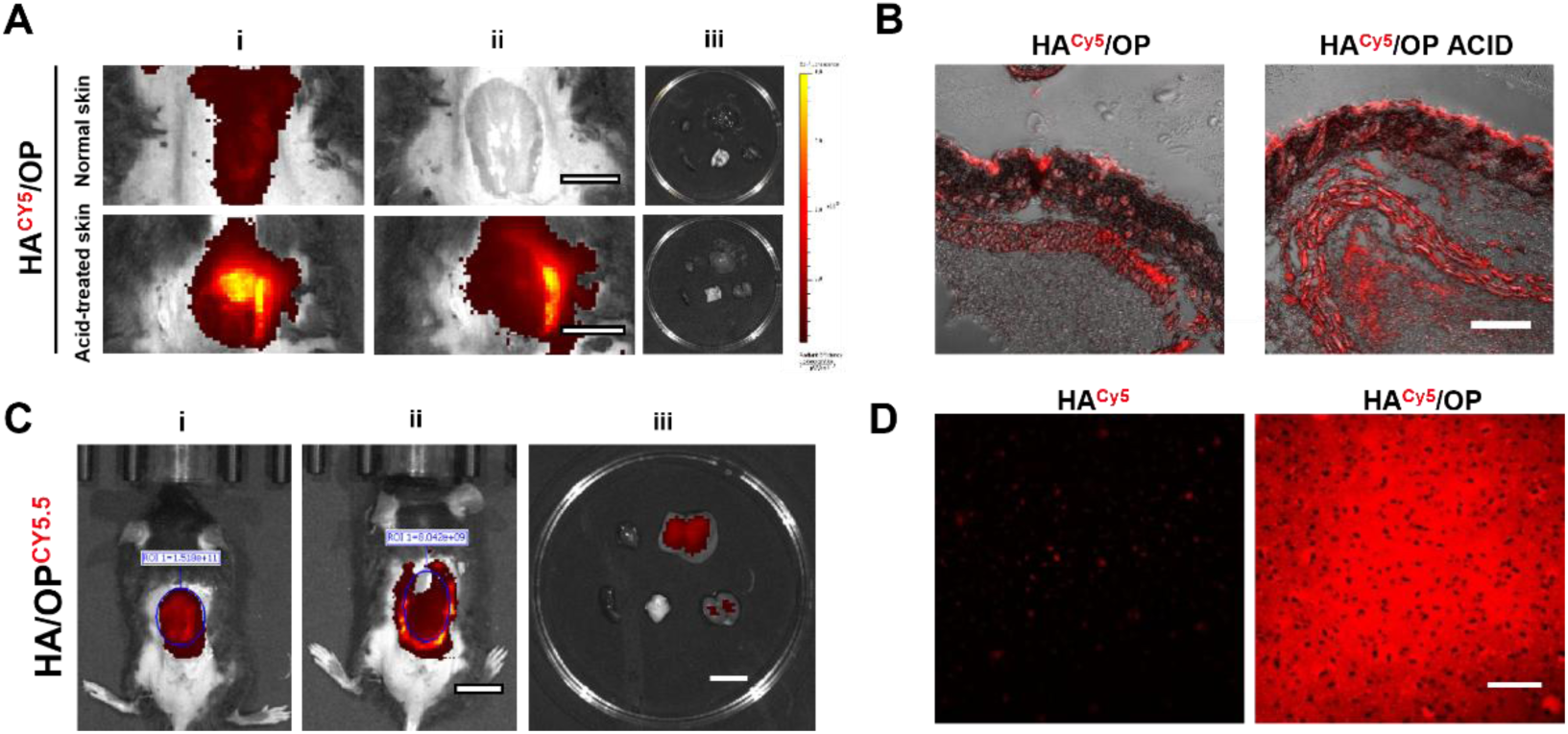
Acid-enhanced skin permeation of HA/OP. (**A**) In vivo cy5-fluorescence imaging of the mouse’s normal skin or acid-pretreated skin topically applied with HA^Cy5^/OP for 4 h (i), the tissue under the skin (ii), and the main organs (iii); (**B**) Representative CLSM images of the skin slices in A. (**C**) In vivo cy5-fluorescence imaging of the mouse’s skin topically applied with HA/OP^Cy5.5^ for 4 h (i), the tissue under the skin (ii), and the main organs (iii); (**D**) CLSM images of SC-removed skin pretreated with HA^Cy5^ and HA^Cy5^/OP for 4h. Scale bars, 10 mm [(A) and (C)], 200 μ m [(B) and (D)].

**Fig. S4.**
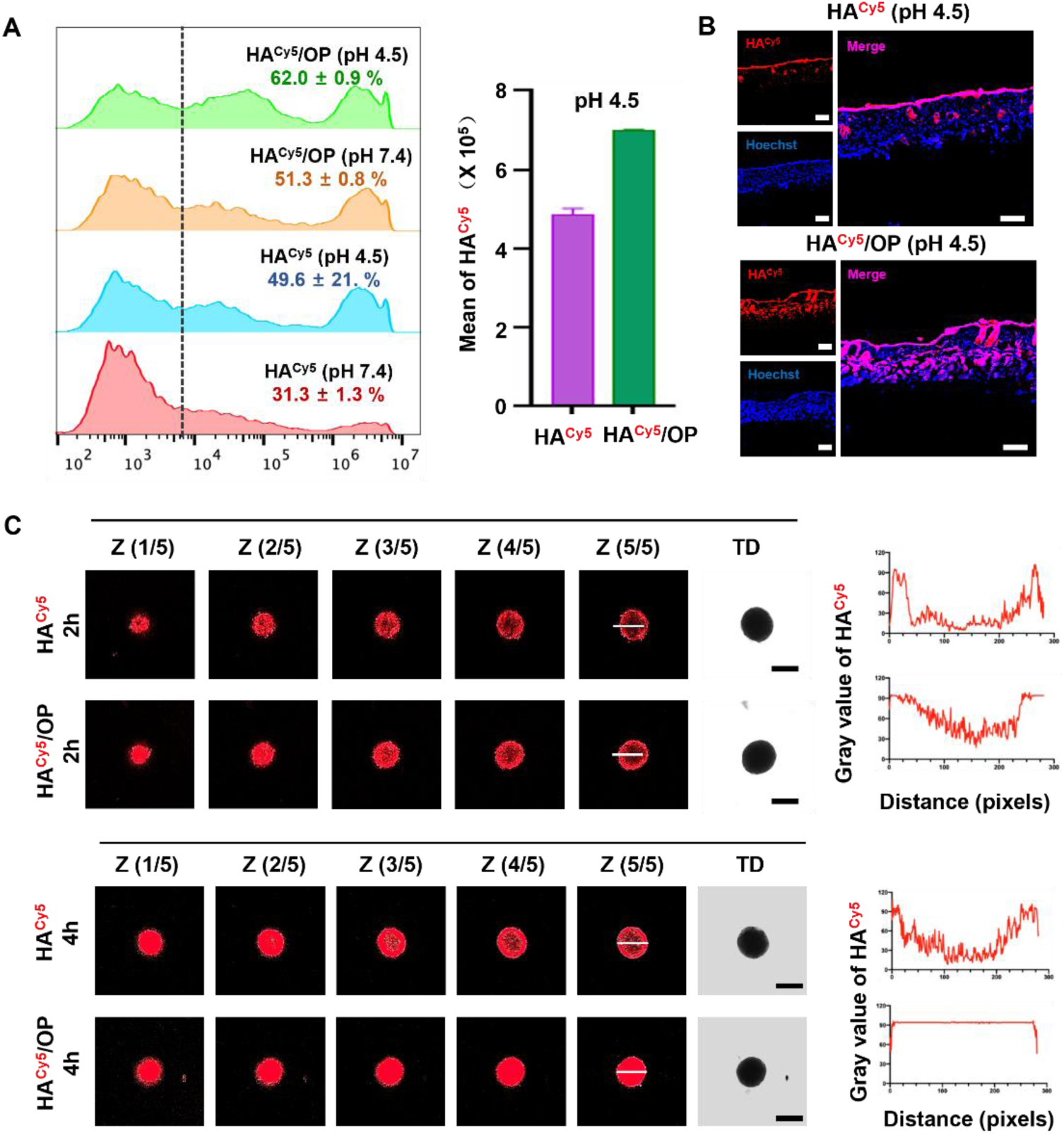
Permeation effect of HA/OP in epidermis and dermis. (A) Cy5 fluorescence detection of the dermis treated with HA^Cy5^ and HA^Cy5^/OP (pH = 7.4 or 4.5) in 4 h were detected using FCM after the stratum corneum removed. (B) Cy5 fluorescence detection of the dermis treated with HA^Cy5^ and HA^Cy5^/OP (pH = 4.5) in 4 h were detected with CLSM after the stratum corneum removed. (C) Cy5 fluorescence detection of HaCaT spheroids treated with HA^Cy5^ and HA^Cy5^/OP in 2 h and 4 h. Scale bars, 50 μm (B), 200 μm [(C) and (D)].

**Fig. S5.**
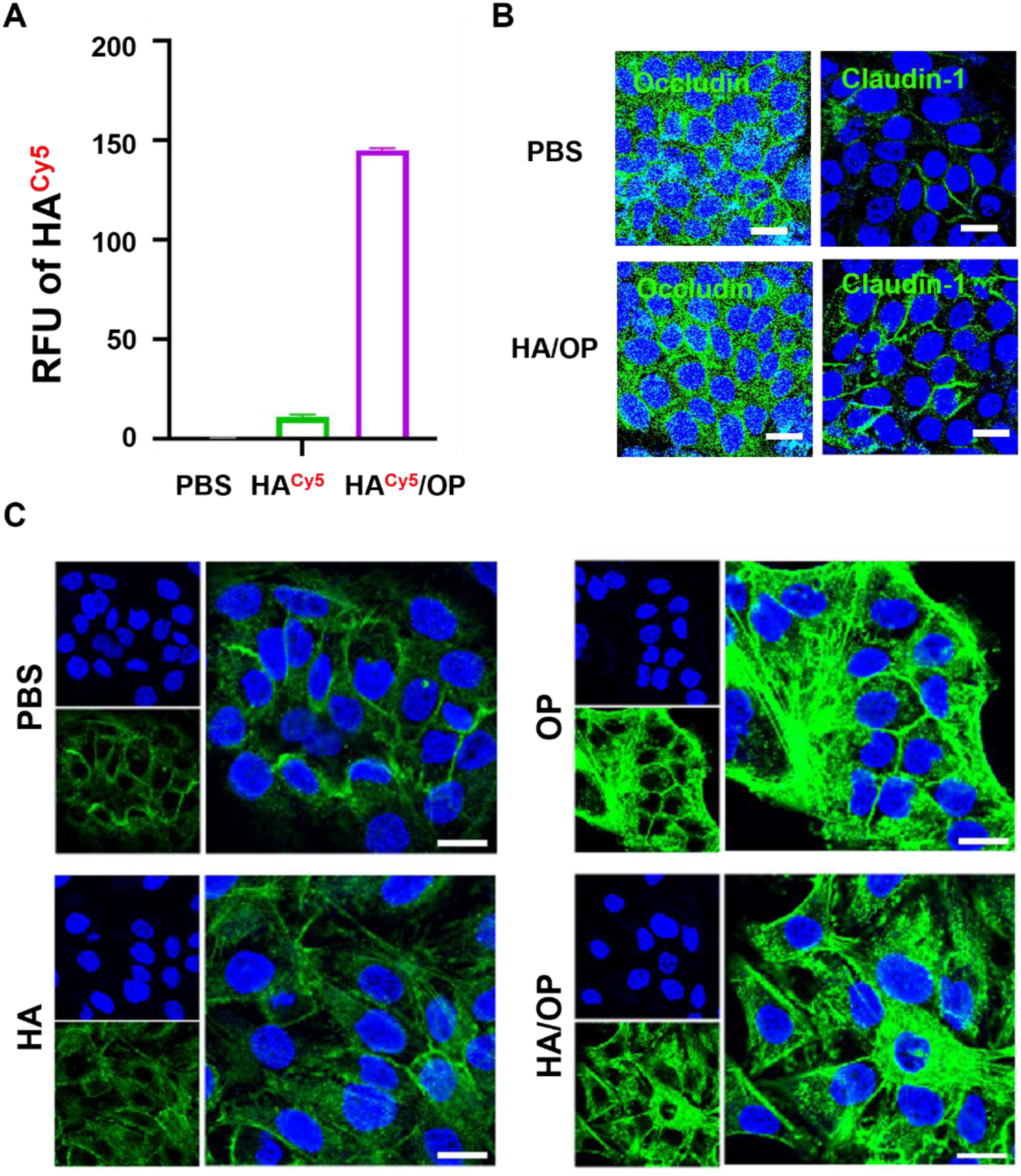
The exchanges of paracellular permeability and tight junctions by HA^Cy5^/OP in HaCaTs. (A) Cy5 fluorescence detection in the outer chamber fluid of the transwell after 4 h incubation with HA^Cy5^/OP. (B) The distribution of Occludin and Claudin-1 after HA/OP was administered to HaCaTs for 4 h by CLSM. (C) Distribution changes of F-actin in HaCaTs after 4 h incubation with HA/OP. Scale bars, 20 μm [(B) and (C)].

**Fig. S6.**
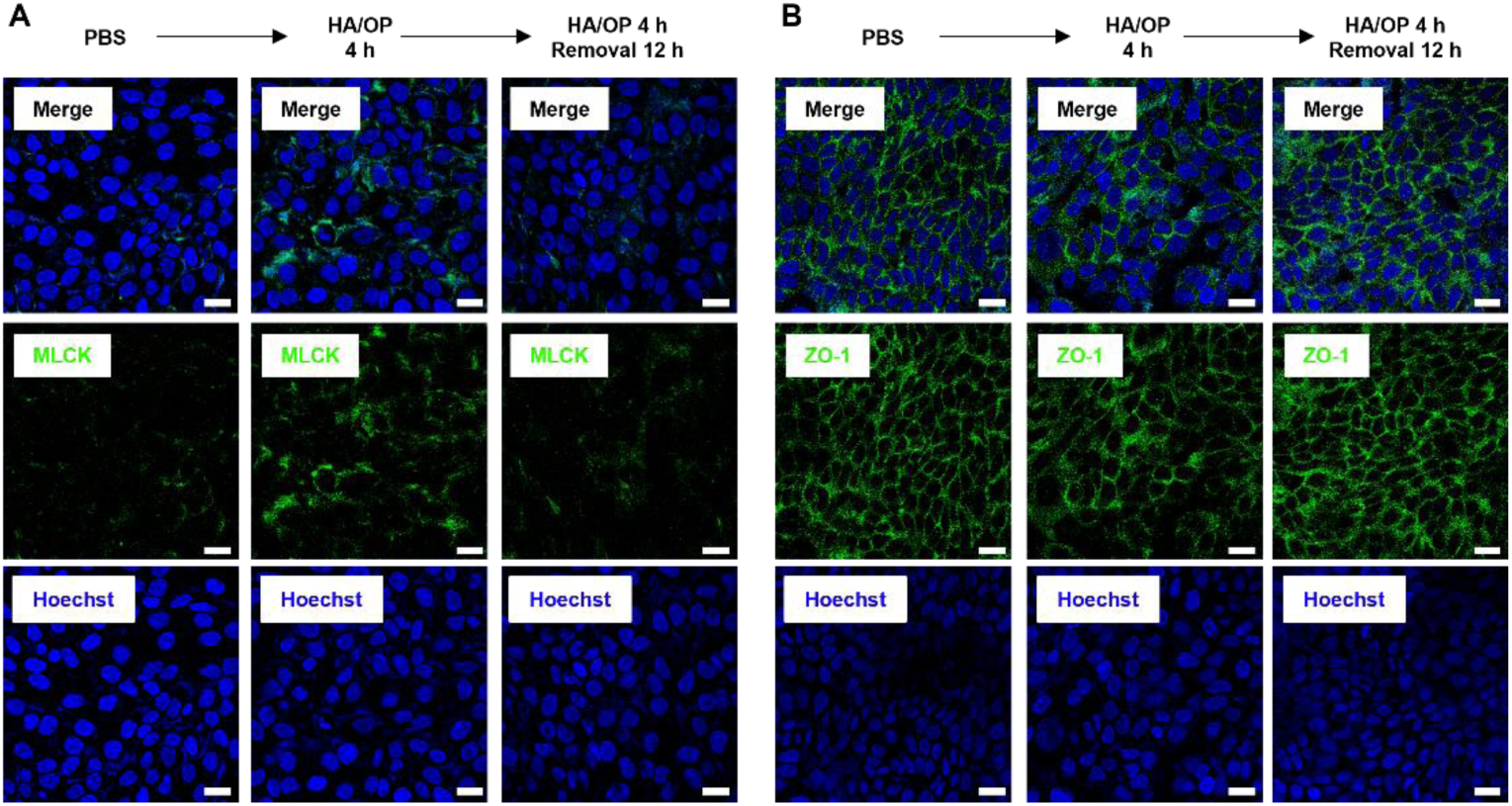
The reversible MLCK-mediated paracellular permeability of HA/OP in HaCaTs. After 4 h of HA/OP administration, HaCaTs were incubated with fresh medium for another 12 h, and the protein amount and distribution of MLCK (A) and ZO-1 (B) were detected by CLSM. Scale bars, 20 μm.

**Fig. S7.**
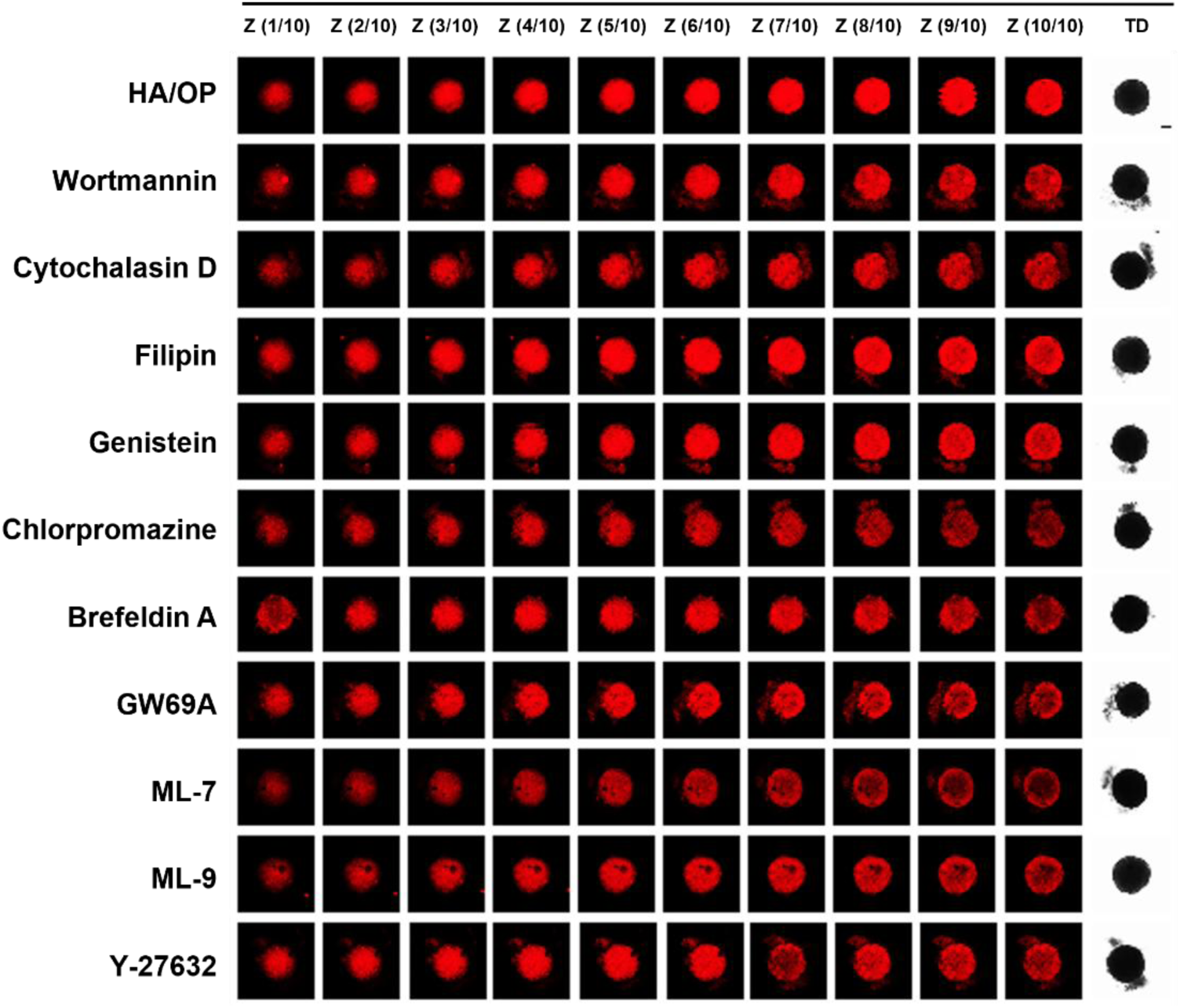
The effects of various inhibitors on the penetration of HA^Cy5^/OP in HaCaT spheroids. After the formation of HaCaT spheroids, the 3D spheroids were pretreated with these inhibitors for 12 h (Wortmannin, 5 μM; Cytochalasin D, 5 μM; Filipin, 10 μM; Genistein, 200 μM; Chlorpromazine, 50 μM; Brefeldin A, 10 μM; GW69A, 20 μM; ML-7, 20 μM; ML-9, 20 μM; Y-27632, 100 μM), and then HA^Cy5^/OP was added to continue the incubation for 4 h. Scale bars, 50 μm.

**Fig. S8.**
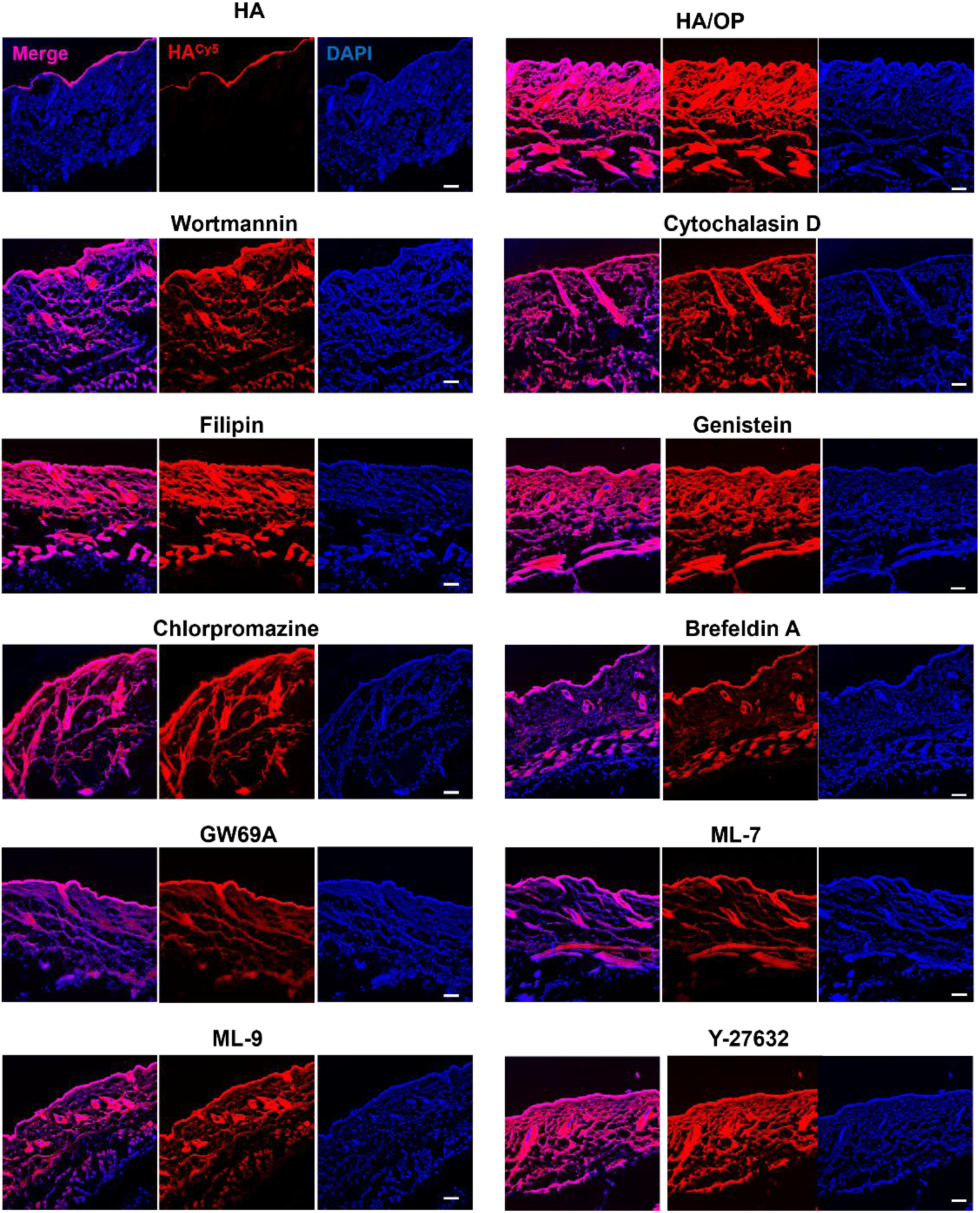
The effects of various inhibitors on the penetration of HA^Cy5^/OP in SC-removed skin. 1 h after subcutaneous injection of ten kinds of inhibitors (Wortmannin, 1.5 mg/kg; Cytochalasin D, 1.5 mg/kg; Filipin, 1.5 mg/kg; Genistein, 5 mg/kg; Chlorpromazine, 2.5 mg/kg; Brefeldin A, 5 mg/kg; GW69A, 2.5 mg/kg; ML-7, 2.5 mg/kg; ML-9, 2.5 mg/kg; Y-27632, 5 mg/kg) and 4 h after administration of HA^Cy5^ or HA^Cy5^/OP, Cy5 fluorescence of the SC-removed skin was detected with CLSM. Scale bars, 50 μm.

**Fig. S9.**
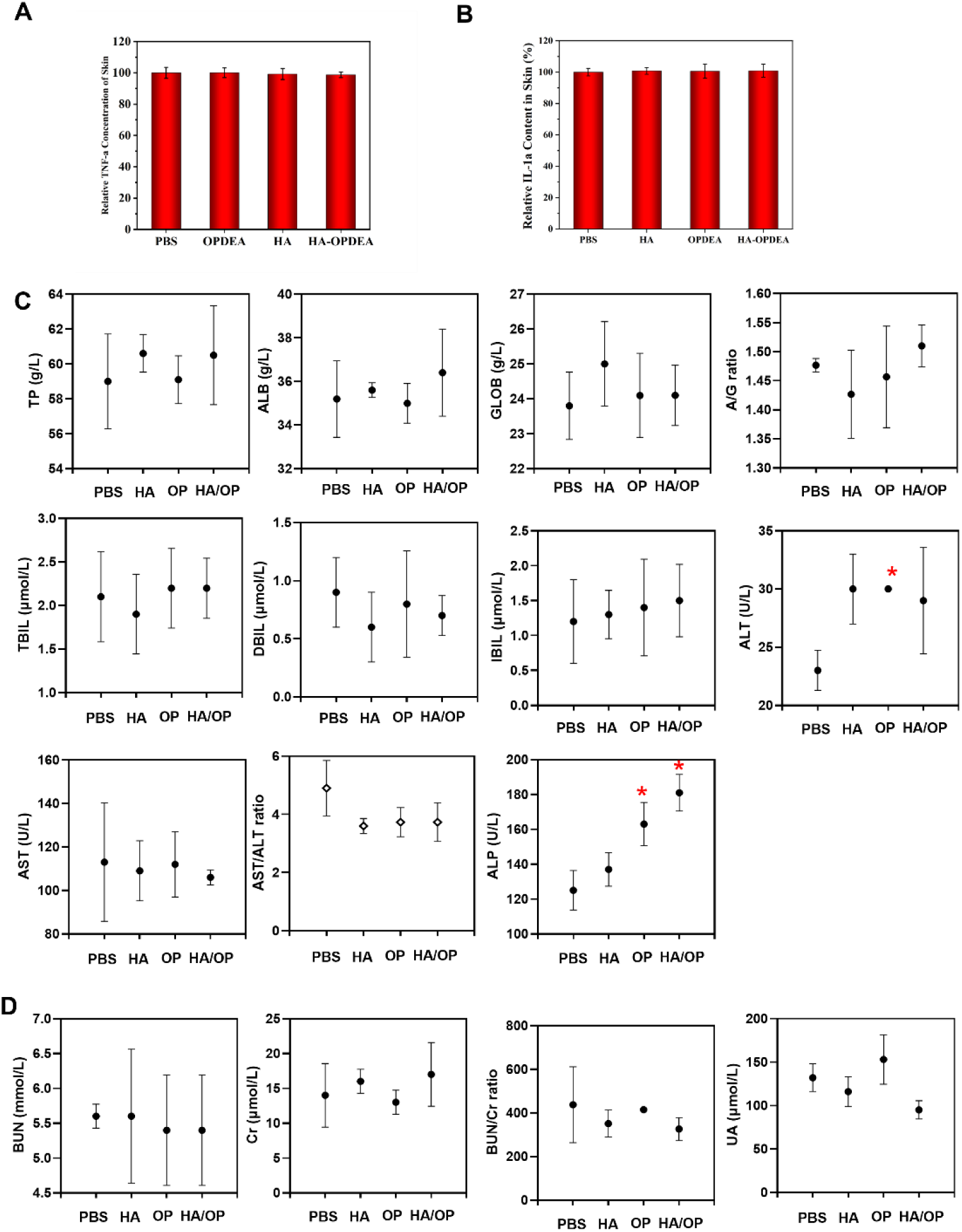
Biocompatibility evaluation of HA/OP. (A) Relative TNF-α concentration in skin pretreated with PBS, OP, HA, and HA/OP for 4 h daily for 15 days. (B) Relative TNF-α concentration in skin pretreated with PBS, OP, HA, and HA/OP for 4 h daily for 15 days. (C) Routine liver function tests (LFTs) in the serum of mice daily topical application of PBS, HA, OP, and HA/OP for 30 consecutive days. (D) Routine renal function tests in the serum of mice daily topical application of PBS, HA, OP, and HA/OP for 30 consecutive days.

